# Perilipin 4 repetitive region forms amyloid fibrils promoted by a genetic expansion found in myopathy patients

**DOI:** 10.1101/2025.06.03.657256

**Authors:** Charbel Choufani, Nicolas Fuggetta, Cyril Moulin, Bayane Sabbagh, Céline Franckhauser, Sebastien Lyonnais, Ana-Andreea Arteni, Sylvie Rouquier, Tinkara Perme, Sophie Pagnotta, Alessandra Ruggieri, Luc Bousset, Alenka Čopič

## Abstract

Lipid-binding amphipathic proteins play key roles in cellular organization, yet their conformational flexibility makes them prone to aggregation. Here we study the aggregation of perilipin 4 (PLIN4), a protein of the lipid droplet (LD)-binding perilipin family. PLIN4 contains an unusually long (∼1000 amino acid, aa) amphipathic helix composed of tandem 33–aa repeats, which folds on the LD surface. A genetic expansion that increases the number of repeats is associated with skeletal muscle dysfunction in patients suffering from a late-onset vacuolar myopathy (MRUPAV). We show that the genetic expansion and the precise sequence of the 33 aa repeats do not significantly alter PLIN4 interaction with LDs in cells. By contrast, purified fragments of the PLIN4 repetitive region form amyloid fibrils as characterized by cryo-EM and atomic force microscopy. Repeat expansion strongly accelerates fibril formation, consistent with enhanced self-assembly driven by the accumulation of identical repeats. PLIN4 aggregation is attenuated by the presence of LDs, indicating that lipid binding modulates the balance between membrane association and self-assembly. Together, these results establish PLIN4 as a previously unrecognized amyloid-forming protein and suggest that repeat expansion enhances amyloidogenic self-assembly, providing a foundation for future studies of its role in degenerative muscle disease.

## Introduction

Many degenerative diseases are associated with protein aggregation (1). Protein aggregates can be amorphous or forming an ordered phase, with the most highly ordered aggregates presenting as amyloid fibrils, where protein units are stacked within a highly stable cross-β fold. Some well-known examples of amyloid forming proteins include α-synuclein, Aβ protein, tau and huntingtin protein (2). Amyloid fibrils represent an alternative conformation, which can be adopted by the wild-type protein and may be promoted by genetic mutations. For example, wild-type α-synuclein is found accumulated in the dense clusters termed Lewy bodies in the brains of patients for which Parkinson’s disease is associated with ageing. However, an early-onset familial form of the disease is linked to genetic mutations in α-synuclein that can promote its aggregation *in vitro* (3). Atomic structures of amyloid fibrils have been determined and show how amino-acid residues are stacked in the cross-β fold (4). Only specific segments of amyloid forming proteins are engaged in the amyloid core, ranging from 15 to 199 amino acids (aa), representing 5 to 100% of the total protein sequence length (5, 6). Amyloid-forming regions often contain repetitive sequences, which can range from stretches of a single aa, such as glutamine (Q), to longer imperfect repeats (7). In huntingtin protein, expansion of the poly-Q tract causes Huntington’s disease, with repeat length correlating with disease severity (8, 9). α-Synuclein contains imperfect 11-aa repeats (10), whereas tau protein has four microtubule-binding repeats of 31-32 aa that differ in their aggregation propensities (11).

A rare progressive distal myopathy that leads to loss of skeletal muscle was first described in 2004 in a large multi-generation family (12) and was termed MRUPAV, for “myopathy with rimmed ubiquitin-positive autophagic vacuolation” (MIM #601846). Distal myopathies are genetically-diverse diseases, often characterized by the presence of protein aggregates in muscle fibers or defects in protein folding or aggregate clearance pathways (13). A. Ruggieri and colleagues have recently shown that this autosomal dominant disease is linked to a large coding expansion in *PLIN4* (14, 15). In muscle biopsies of patients carrying the *PLIN4* expansion, perilipin 4 (PLIN4) is the most abundant protein that accumulates in protein-dense enlarged vacuoles and is also observed in subsarcolemmal aggregates. These aggregates are positive for ubiquitin and autophagy markers p62 and NBR1, suggesting activation of the aggrephagy pathway for aggregate clearance (16, 17). PLIN4 contains an unusually long central repetitive region, composed of about 30 highly conserved 33-aa repeats, which bear similarity to the 11-aa repeats of α-synuclein (10, 18). The coding expansion identified in MRUPAV patients results in additional 33-aa repeats in the repetitive region (15). The high repetitiveness of the corresponding DNA region and lack of any nucleotide change in the genetic expansion explains why this mutation was so difficult to identify. Furthermore, MRUPAV patients exhibit clinical symptoms similar to other vacuolar myopathies. Nevertheless, additional families with muscular degeneration linked to *PLIN4* expansion have recently been reported, with one family carrying an even longer expansion, linked to an earlier disease onset and more proximal muscle involvement (19–21).

PLIN4 belongs to the family of perilipins, which are the most abundant proteins on the surface of lipid droplets (LDs) – cellular organelles dedicated to the storage of energy in the form of neutral lipids (22, 23). LDs are prominent in myofibrils and provide energy for muscle contraction. Their number and properties can vary in disease states, such as type II diabetes, or with training (24, 25). PLIN4 is highly expressed in skeletal muscle and in adipocyte cells (26, 27). Whereas perilipins are generally considered as regulators of LD metabolism, the molecular function of PLIN4 remains enigmatic. Recent evidence suggests that PLIN4 is important for regulation of LD stability, in particular during increased LD production (28). Furthermore, analysis of whole-exome sequencing data from a large cohort shows that loss-of-function mutations in *PLIN4* correlate with unfavorable body fat distribution and metabolic profiles in humans (22, 29). These studies point to an important role of PLIN4 in the regulation of cellular lipid metabolism.

We have previously shown that the repetitive region of PLIN4 directly interacts with the LD surface and is required for PLIN4 targeting in cells, where PLIN4 can be found both in the cytosol and on LDs. The repetitive region is disordered in solution and it folds into an amphipathic helix in contact with LDs with which it interacts in a length-dependent manner (18, 30). Amphipathic helices are common in LD-binding proteins. However, PLIN4 is unique in the size and highly repetitive nature of its amphipathic helix sequence. Interestingly, the much shorter and imperfectly-repetitive region of α-synuclein, which forms amyloid fibrils, has also been shown to target LDs (18, 31, 32). LDs have been observed in vicinity of α-synuclein aggregates and are generally implicated in protein quality control (33–35).

Here, we explore the molecular mechanism of the *PLIN4*-linked MRUPAV myopathy. We show that the increased number of the 33-aa repeats in PLIN4, introduced by the genetic expansion, or their exact sequence do not significantly impact the interaction of PLIN4 with LDs in cells. However, the purified repetitive region of PLIN4 can form amyloid fibrils, and the kinetics of fibril formation is highly promoted by the MRUPAV mutation, which leads to multiplication of identical 33-aa repeats. By contrast, the presence of LDs decreases the rate of fibril formation in vitro. The genetic expansion carried by the MRUPAV patients may therefore represent a toxic gain-of-function mutation that promotes aggregation of PLIN4 in skeletal muscle tissue.

## Results

### The genetically-encoded expansion of PLIN4 does not affect its interaction with LDs

PLIN4 (Uniprot Q96Q06) contains a long repetitive region composed of highly similar 33-aa repeats, flanked by a disordered N-terminal region and a folded C-terminal domain. The repetitive region is reported to contain 29 to 31 repeats in the wild-type protein, and is expanded by 9 or more repeats in MRUPAV patients: in this study, we compare the Uniprot reference sequence with the mutant sequence described by Ruggieri et al. (15), which contains 9 additional 33-aa repeats (Fig. 1A, B). The repetitive region of PLIN4 is primarily responsible for PLIN4 interaction with LDs, folding into an amphipathic helix in contact with LDs. Heterologous expression of PLIN4 fragments containing different number of repeats has shown that while the affinity for LDs per individual repeat is low, it increases with the number of repeats (18, 30). We first asked whether the genetic expansion affects the localization of full-length PLIN4 to LDs. In cultures of myoblasts and myotubes obtained from the patients’ muscle biopsies, PLIN4 expression is not detected (A Ruggieri, unpublished data). We therefore developed heterologous cellular models to compare the properties of wild-type and mutant PLIN4.

**Fig. 1.**
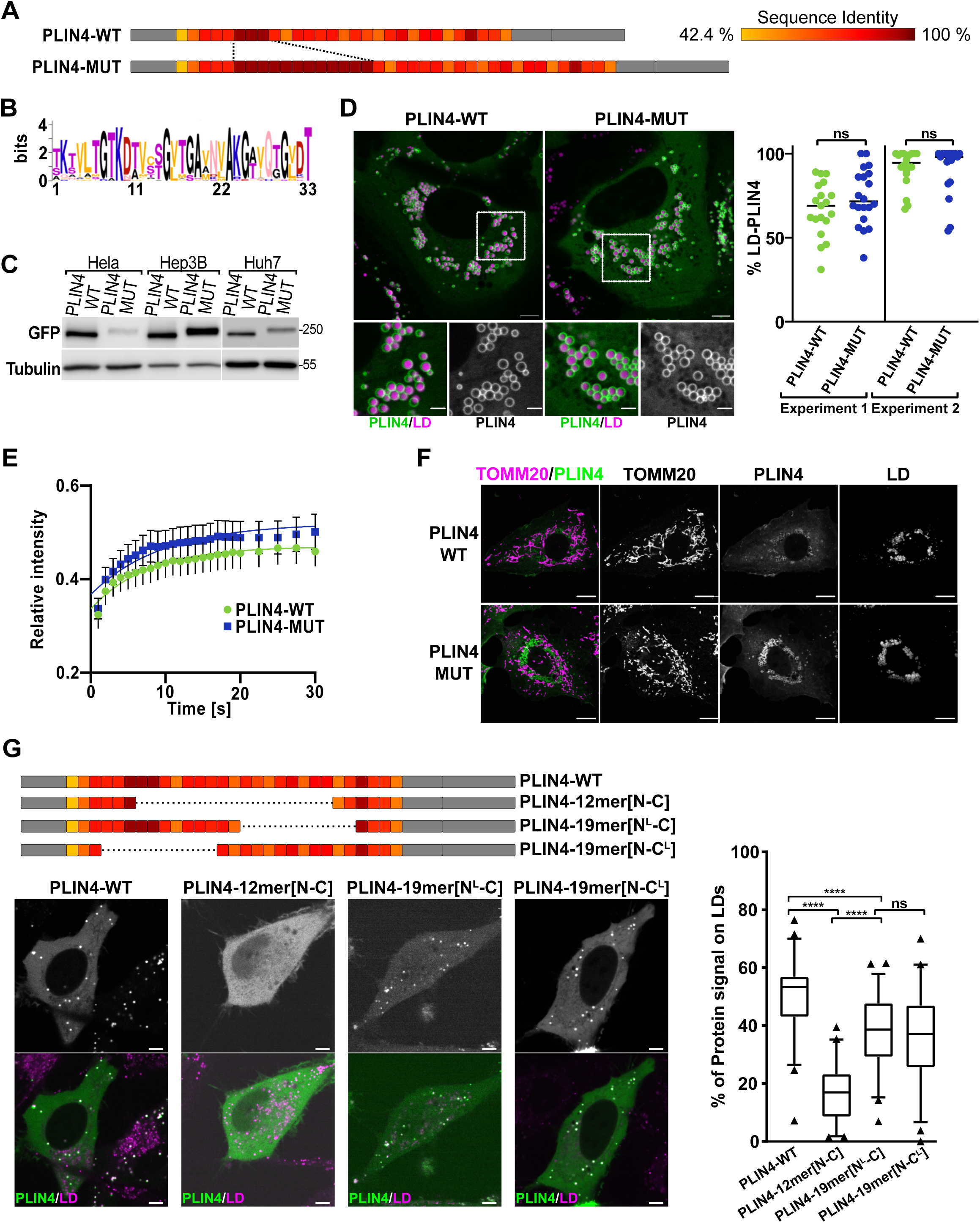
The effect of genetic expansion in PLIN4 on its interaction with LDs. (A)Schematic representation of wild-type (WT) (Uniprot Q96Q06) and mutant (Mut) (15) forms of human PLIN4. The repetitive region, composed of 29 x 33-aa repeats in PLIN4-WT and 38 x 33-aa repeats in PLIN4-Mut is shown in color using a heat map that represents the degree of sequence identity between the repeats (see legend on the right and Table S1). (B)Weblogo showing conservation between the 29 33-aa repeats present in WT PLIN4. (C)Western blot analysis of PLIN4-GFP levels in total cells lysates from three different cell lines (HeLa, Huh7 and Hep3B) expressing PLIN4-WT-GFP or PLIN4-Mut-GFP after doxycylin (Dox) induction. Tubulin was used as a loading control. (D)Localization of WT and Mut PLIN4-GFP in Huh7 cells after Dox induction as in C), in media containing oleic acid (OA). LDs were stained with Lipi-Blue prior to live imaging by confocal microscopy. Bottom panels show a zoom-in of the selected areas. Scale bars: 5 μm and 2 μm for the zoom-in. Graphs show percent of LDs positive for PLIN4 in individual cells quantified in two independent experiments (exp. 1, n=19 for WT and n=20 for Mut cell line; exp. 2, n=20 for WT and n=19 for Mut cell line). Samples in the same experiment were compared using Mann-Whitney test, ns, nonsignificant (P>0.05). (E)FRAP analysis of PLIN4-GFP signal (WT or Mut) on LDs in Huh7 cells grown as in D). Data points represent mean of 20 recovery measurements ± SEM. (F)Analysis of mitochondria in Huh7 cells expressing PLIN4-GFP WT or Mut by IF against mitochondrial protein TOMM20. PLIN4-GFP was detected with antibodies against PLIN4 and LDs were stained with Lipi-Blue. Scale bars: 10 μm. (G) Localization of PLIN4-GFP constructs with deletions in the repetitive region, transiently expressed in HeLa cells. A construct containing 12 repeats (PLIN4-12mer[N-C]) and two constructs containing 19 repeats, with deletions of different regions, as indicated in the diagram, were compared with PLIN4-WT (29 repeats). Quantification shows % of protein signal on LDs per cell, n=50 for each construct. Samples were compared using a pairwise t-test, ****P <0.0001.

We created several cell lines expressing wild-type (WT) or mutant (MUT) form of full-length PLIN4, tagged with GFP at the C-terminus, from an inducible promotor using the sleeping-beauty system.

In two hepatocyte cell lines, the two forms of PLIN4 reached similar protein levels upon doxycycline induction, whereas in HeLa cells, the levels of PLIN4-MUT were consistently lower than those of PLIN4-WT (Fig. 1C). We therefore used the hepatocyte cell lines to compare the affinity of PLIN4-WT and MUT for LDs. Both proteins localized to the LD surface as well as to the cytosol, showing similar distribution profiles (Fig. 1D and Fig. S1A). We also observed similar fluorescence recovery after photobleaching (FRAP) for the two proteins (Fig. 1E and Fig. S1B). These results suggest that the expansion of the PLIN4 repetitive region does not strongly impact PLIN4 interaction with LDs. Furthermore, we did not observe significant differences in the morphology of the mitochondrial network nor in the number of LDs (Fig. 1F and Fig. S1C).

To further explore how the number and aa sequence of the 33-aa repeats in the context of full-length PLIN4 impact its interaction with LDs, we prepared GFP-tagged PLIN4 truncation mutants with deletions in different parts of the repetitive region and analyzed their binding to LDs in transiently-transfected HeLa cells. Decreasing the total number of repeats from 29 to 12 strongly decreased PLIN4 association with LDs, whereas the difference between the 29 and the 19-repeat construct was smaller. Two 19-repeat PLIN4 mutants with deletions of different parts of the repetitive region displayed very similar LD-localization patterns (Fig. 1G), suggesting that variations in the aa sequence of individual PLIN4 repeats do not strongly affect the binding to LDs, in agreement with the analysis of the repetitive region alone (18).

To further evaluate the effect of the genetic expansion on PLIN4, we turned to *in vitro* experiments with purified proteins. The nature of PLIN4 (combination of long disordered and repetitive regions and a folded C-terminal domain) makes purification of the full-length protein extremely challenging. We therefore focused on the repetitive region alone in these experiments. We first compared the biochemical properties of two PLIN4 fragments of the same length (containing 12 x 33-aa repeats) from two regions of the WT protein: 12mer[260:] starts at aa 260 and includes the region that is affected by the genetic expansion; 12mer[524:] includes a down-stream region (Fig. 2A). Both fragments were purified in the same manner, as previously established for 12mer[524:] (Fig. S2A) (18). We used circular dichroism (CD) to compare their secondary structures. In buffer, the two constructs display spectra typical for unstructured proteins, whereas they show a strong shift to α-helical structure in the presence of 50% trifluoroethanol (TFE), as previously shown for 12mer[524:] (Fig. 2B) (18). Furthermore, both fragments are highly susceptible to limited proteolysis by subtilisin, indicative of high exposure of peptide bonds all along the sequence, i.e., a lack of stable structure. By contrast, the two fragments become more resistant to proteolysis by the addition of liposomes, indicating their folding on the liposome surface (Fig. 2C) (36). Of note, these liposomes contain diphytanoyl phospholipids with branched acyl chains to mimic packing defects at the surface of LDs (18, 28). Overall, these cellular and biochemical data suggest that the genetically-encoded expansion in the PLIN4 repetitive region present in the MRUPAV patients have no significant impact on the interaction of PLIN4 with LDs.

**Fig. 2.**
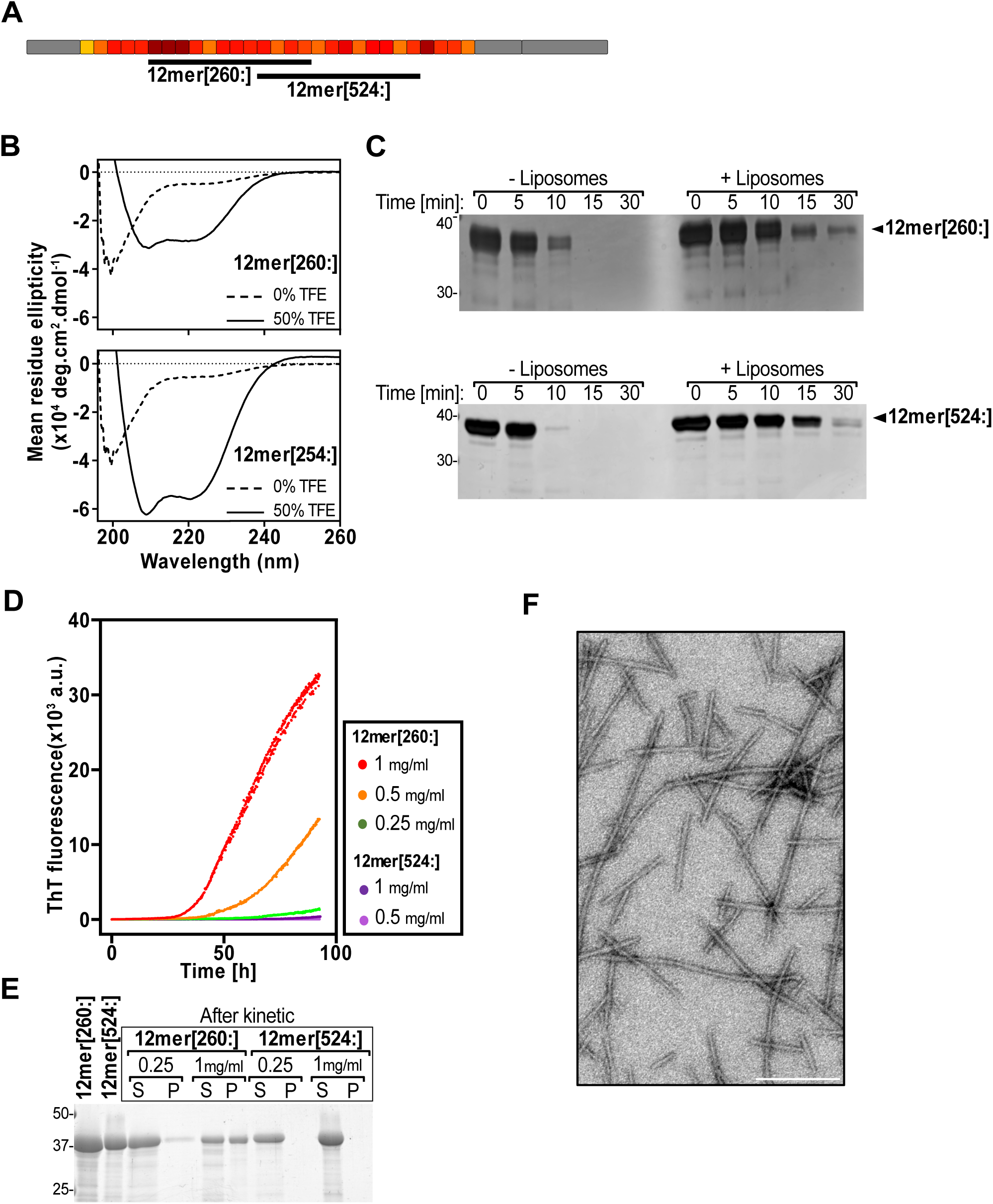
PLIN4 repetitive regions is prone to formation of fibrils. (A)Schematic representation of PLIN4-WT (Uniprot Q96Q06) with the repetitive region highlighted in color, showing the positions of the constructs 12mer[260:], 12mer[524:]. (B)CD spectra of 12mer[260:] (above) and 12mer[524:] (below) (13 µM and 14 µM, respectively) in buffer or in the presence of 50% TFE. (C)Limited proteolysis of 12mer[260:] (above) and 12mer[524:] (below) (0.1 mg/ml for both proteins) with 0.2 µg/mL subtilisin, in the presence or absence of liposomes. Proteins were analyzed by SDS-PAGE and stained with Silver stain. Data shown is representative of 2-3 independent experiments. (D)Thioflavin T (ThT) assay to monitor the aggregation kinetics over 90 hours (h) of 12mer[260:] at 1 mg/mL (red), 0.5 mg/mL (orange) and 0.25 mg/mL (green) and 12mer[524:] at 1 mg/mL (purple). (E)Pelleting assay (left) of 12mer[260:] and 12mer[524:] at 0.25 mg/mL and 1 mg/mL before and after the ThT assay carried out for 100 h (right). Pellets and supernatants after centrifugation at 100 000 g were analyzed by SDS PAGE and stained with GelcodeBlue. (F)Fibrils obtained by aggregation of 12mer[260:] at 0.5 mg/ml in a ThT assay were visualized by negative stain EM. Scale bar: 0.2 µm.

### Fragments of PLIN4 can form amyloid fibrils

In skeletal muscle fibers of MRUPAV patients, PLIN4 accumulates and co-localizes with markers of the aggrephagy pathway, suggesting that it may be forming aggregates (15). Therefore, we asked whether the repeat sequences might be susceptible to adopting alternative conformations.

We first noticed the formation of regular filamentous structures in a concentrated solution of a longer PLIN4 fragment 20mer[260:]. kept at room temperature for 1 week before observation by negative stain electron microscopy (EM) (Fig. S2B). This observation prompted us to evaluate the formation of PLIN4 filaments under controlled conditions. We incubated the two afore-mentioned 12mer fragments at different concentrations at 37°C under agitation in the presence of ThioflavinT (ThT), a fluorescent compound whose intrinsic fluorescence drastically increases after intercalation into protein fibrils (amyloids) (37). Strikingly, we observed a large and concentration-dependent increase in ThT fluorescence over the course of several days with 12mer[260:], whereas 12mer[524:] showed almost no increase in ThT fluorescence even at the highest concentration (Fig. 2D). We also assessed the formation of fibrils using a pelleting assay and observed a good correlation with the ThT assay (Fig. 2E): before incubation, both proteins remained in the supernatant, whereas after incubation at 37°C for 100 h, ∼10% of 12mer[260:] at 0.25 mg/ml and ∼50% of 12mer[260:] at 1 mg/ml were found in the pellet fraction. 12mer[524:] at 1 mg/ml remained soluble. Analysis of protein samples after ThT incubation by negative-stain EM consistently revealed abundant and uniform fibrils formed by 12mer[260:] (Fig. 2F). For 12mer[524:] we could observe fibrils in only one experiment, following a prolonged incubation at very high protein concentration (2 mg/ml, Fig. S2C). Together, these results suggest that the repetitive region of wild-type PLIN4 is prone to fibril formation, especially the region comprising the repeats that are multiplied in MRUPAV patients.

To determine the characteristics of the 12mer[260:] fibrils, we performed cryo-EM and atomic force microscopy (AFM) in liquid (Fig. 3). The cryo-EM analysis of vitrified PLIN4 fibrils has allowed extraction of 20 000 filament segments. Segments were boxed into 788 x 788 Å squares and aligned with Relion4 helical module (Fig. 3A). Two main classes were identified, thin and thick filaments. The pattern analysis of filaments by Fourier transform unequivocally shows that PLIN4 fibrils are amyloid in nature, as revealed by the 4.77 Å amyloid spacing, which is the signature of a cross-β pattern (Fig. 3B,C) (2). However, despite the high quality of our cryo-EM data, averaging revealed heterogeneity within the fibrils, suggesting that there might be different modes of stacking between the repeats of the 12mer[260:] PLIN4 fragment during amyloid growth. Observation of 12mer[260:] fibrils by AFM revealed amyloid fibrils that were heterogeneous in length, with a left-handed periodic twist and a maximum height between 7 and 8 nm (Fig. 3D). Periodicity analysis by AFM (38) showed an average distribution around 60 nm with significant variations, in agreement with the heterogeneity noted by CryoEM (Fig. 3E,F). In addition, the height profiles show atypical variations suggesting gaps in the fibril structure (Fig. 3E). Multiple scanning on single fibrils led to fibril disruption, eventually generating destructed ends with short sections around 3-4 nm in diameter that did not show two identifiable fibrils, suggesting that this is the diameter of a single PLIN4-12mer[260:] fibril (Fig. 3G).

**Fig. 3.**
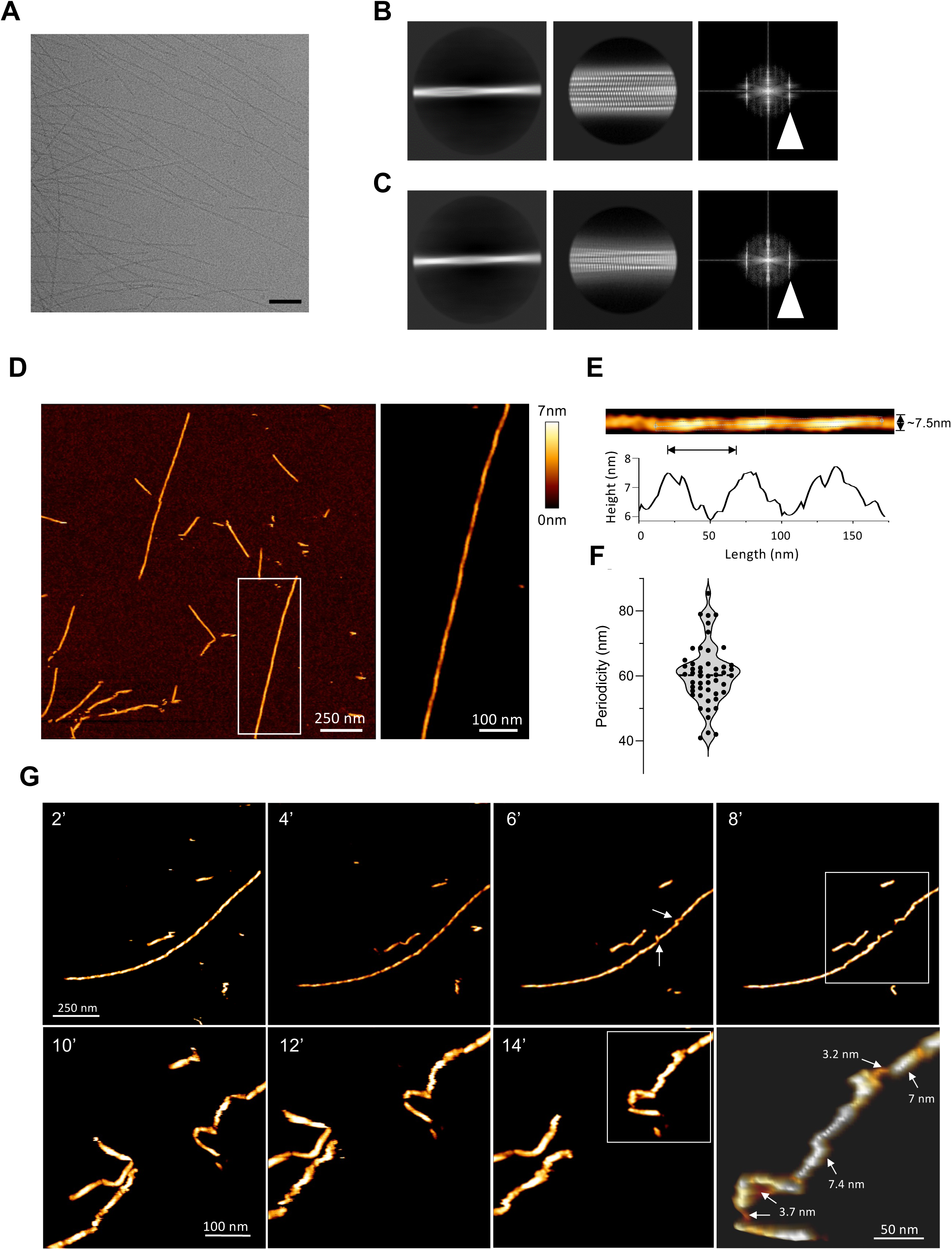
PLIN4 fibrils are amyloid in nature. (A) Fibrils formed from Plin4-12mer[260:] observed by cryo-EM. Scale bar: 100 nm. (B and C) 2D classification of Plin4-12mer[260:] using Relion. Representative 2D classes using 1350 Å box size binned to 3.85 Å/pixel, which identifies two polymorphs with a thick diameter (B, left panel) and thin diameter (C, left panel). Middle panels: 2D classification of the thick and thin polymorph with un-binned images (0.77 Å/pixel) revealing a typical amyloid spacing. Right panels: Fourrier transform of the 2D class shows a strong reflection at 4.79 Å for both thick and thin polymorphs. (D)Plin4-12mer[260:] fibrils formed from Plin4-12mer[260:] were deposited onto freshly cleaved mica and imaged in buffer in PeakForce tapping mode AFM (scale bar: 250 nm). The right panel is a high-resolution magnification of the fibril in the white box, highlighting the twist pattern (scale bar: 100 nm). (E)Straightened fibril and its corresponding height profile, showing a periodicity in the height profile of a single fibril. The approximal length of the repeating unit between two height maxima (periodicity) is indicated. (F) Distribution of periodicities measured for about 15 different fibrils, as shown in (E); each dot is the measurement of one periodicity (n=50). (G) Multiple scans were recorded every 2 min on a single fibril at low setpoint (300 pN), resulting in fibril breakage (scale bar: 250 nm). In the 3rd upper panel, recorded at 6 min, the arrows indicate punctual distortion of the fibrils before breaking (boxed area at 8 min). The lower panels show a 2.5x magnification (scale bar: 100 nm) of the same area that is shown in the box at 8 min containing the breakage site, at 10, 12 and 14 min. The right-most bottom image shows a 2x magnification of the broken fibril in the white box at 14 min, with arrows indicating height profiles of the visible structures (scale bar: 50 nm).

In summary, our structural analysis shows that the repetitive region of PLIN4 can form amyloid fibrils with variable periodicity and structural heterogeneity, and raises questions regarding how the 33-aa repeats of PLIN4 interact within the amyloid fibril.

### Influence of PLIN4 repeat sequence and number on fibril formation

Next, we asked if any feature of the extended version of PLIN4, present in the muscle fibrils of MRUPAV patients, would increase the propensity of the protein towards amyloid formation. We first investigated how the exact sequence of its repeats influences PLIN4 fibrillation. Overall, the repeats display unique aa composition, with some aa highly over-represented (valine, glycine, threonine), under-represented (arginine, glutamic acid, phenylalanine) or completely absent (tryptophan, tyrosine, histidine) (30) (Fig. 1B). We used amyloid prediction algorithms ArchCandy2, Pasta and Tango, included in the TAPASS pipeline (39–42), to determine the size and position of amyloidogenic regions (ARs) within the PLIN4 sequence. The majority of predicted ARs were identified within the repetitive region. Their median length was 27 aa, and their position matched the definition of the reading frame of the repeats, *i.e.,* they mapped within the individual repeats (Fig. S3 and Table S1), meaning that these segments could form the amyloid core of PLIN4 fibrils. We then used ArchCandy 2 to predict the propensity of individual repeats to form amyloids (Table S1). The prediction is plotted as a heatmap for individual repeats (below the heatmap of repeat sequence identity) and shows that amyloid propensity varies between individual repeats (Fig. 4A). To experimentally evaluate these results, we analyzed fibril formation in the ThT assay for three 33-aa synthetic peptides corresponding to three PLIN4 repeats with different amyloidogenic propensities (intermediate, high and low), 1mer[260:] (repeat r6 that is multiplied in patients), 1mer[656:] (r18) and 1mer[755:] (r21), respectively (Fig. 4A). We observed rapid fibril formation for 1mer[260:] with a very short lag phase, and the rate of fibril formation was even higher for 1mer[656:] at similar peptide concentrations. By contrast, 1mer[755:] did not show any increase in ThT signal even at the highest concentration (1 mg/ml) (Fig. 4B).

**Fig. 4.**
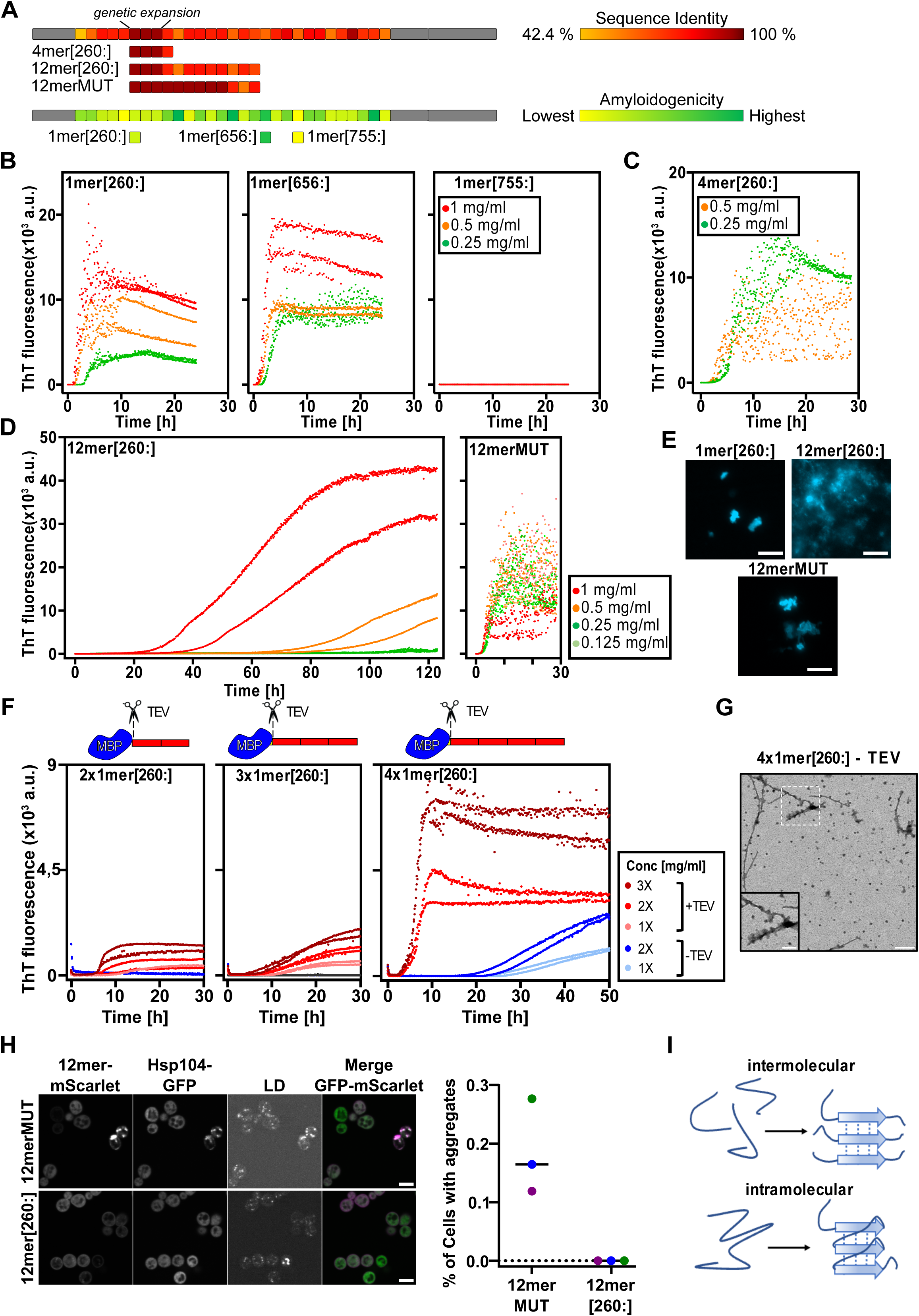
The influence of PLIN4 repeat sequence and number of repeats on aggregation. (A) Diagram of PLIN4 showing %identity between individual 33-aa repeats compared to repeat 6 (r6 = 1mer[260:]), which is multiplied in patients, as a heat map score. Color scale: from 42,4% identity in yellow (minimum %identity) to 100% identity score in red (see Table S1). The expanded region is composed of 3 strictly identical repeats. Represented below: 4mer and 12mer constructs from the WT sequence (containing 3 identical repeats), and the 12mer[260:]MUT construct with 9 identical repeats. Bottom diagram: Heat map scoring of the amyloidogenic propensity of individual 33-aa repeats, determined by ArchCandy (see Table S1). Positions of the three 33-aa peptides evaluated experimentally, 1mer[260:], 1mer[642] and 1mer[741], are shown below. (B) Aggregation kinetics in the ThT assay of peptides representing single 33-aa repeats, 1mer[260:], 1mer[642:] and 1mer[741:], at 1 mg/mL (red), 0.5 mg/mL (orange) and 0.25 mg/mL (green). (C) Aggregation kinetics in the ThT assay of 4mer[260:] at 0.5 mg/mL (orange) and 0.25 mg/mL (green). (D) Aggregation kinetics in the ThT assay of 12mer[260:], compared to 12merMUT at 1 mg/mL (red), 0.5 mg/mL (orange), 0.25 mg/mL (deep green) and 0.125mg/mL (light green). (E) After aggregation in the presence of ThT for 100 h at 37°C, samples of 1mer[260:] (0.5 mg/mL), 12mer[260:] (0.25 mg/mL) and 12merMUT (0.25 mg/mL) were observed by widefield fluorescent microscopy to detect ThT fluorescence. Contrasts in the three images were adjusted to the same levels. Scale bar: 10 µm. (F) ThT aggregation kinetics of MBP constructs containing 2, 3 or 4 tandem identical repeats of the 33-aa sequence 1mer[260:], in the presence (pink to red) or absence (blue) of TEV protease. Two or three different concentrations (1X, 2X or 3X) were tested for each protein, with some variability in the 1X concentration: MBP-2x1mer[260:] = 0.4 mg/mL; MBP-3x1mer[260:] = 0.5 mg/mL; MBP-4x1mer[260:] = 0.55 mg/mL. Proteins were tested immediately after purification without freeze-thawing to minimize protein degradation. (G) Reaction mixtures after ThT aggregation were analyzed by negative stain EM. Fibrils were detected only in the MBP-4x1mer[260:] samples + TEV and – TEV (see Fig. S6A). Scale bar: 1 µm or 0.5 µm for the magnified area (2 x zoom). (H) Localization of 12mer[260:] and 12merMUT-mScarlet fusion proteins in budding yeast with endogenously-tagged Hsp104-GFP. Cells were grown for 18h in selective medium at 30°C prior to observation, and LDs were stained with AutoDOT. Scale bar: 5 µm. Graph shows quantification of the percent of cells with observable PLIN4-12mer aggregates that colocalize with Hsp104-GFP, as shown in (C). Each color symbol on the graph represents an independent experiment. Cell counts for were 12merMUT: Blue, n=7298, Green, n=20958, and Purple, n=10092, and for 12mer[260:]: Blue, n=8753, Green, n=32275, and Purple, n=7569. The value for 12mer[260:] was 0 in all experiments. (I) Model of intermolecular vs. intramolecular amyloid stacking.

We also analyzed the formation of fibrils by longer fragments containing 4 repeats from different parts of the PLIN4 repetitive region (Fig. S4A). Surprisingly, the fragment 4mer[260:], comprising PLIN4 region 260-523 (Fig. 4A), aggregated with a similar kinetics as the single repeat 1mer[260:] at high protein concentration (0.5 mg/ml, Fig. 4C). This was despite its 4-times higher length, which we would expect to slow down the aggregation kinetics due to higher flexibility of the polypeptide chain, as was the case for 12mer[260:] (Fig. 4D). However, the aggregation kinetics of 4mer[260:] was largely concentration-independent and occurred even at 20x lower protein concentration (0.03 mg/ml, Fig. S3B). Of note, unlike all other purified PLIN4 fragments used in these experiments, 4mer[260:] contains a T7 tag of 13 aa at its N-terminal, which was introduced during cloning (see Table S2). This tag likely slightly inhibits the aggregation of this PLIN4 fragment (see the behavior of MBP-fusion constructs described in the next section). Other 4mer fragments (4mer[326:], 4mer[557:], and 4mer[689:]) aggregated with different kinetics that were more or less concentration dependent, with 4mer[557:] showing the most similar behavior to 4mer[260:], 4mer[326:] aggregating more slowly, and 4mer[689:] requiring a much higher concentration for robust aggregation (Fig. S4B-E). This data is in agreement with the prediction that individual repeats have different amyloidogenic propensities and shows that combination of different repeats can lead to variable outcomes.

### Identical repeats in expanded PLIN4 strongly promote fibrillation

A unique feature of the region of the PLIN4 sequence that is expanded in majority of MRUPAV patients is that it contains identical repeats (r6-r8, aa260-358); the mutant PLIN4 proteins in the muscle tissue of these patients therefore contain 12 or more identical repeats as compared to 3 in wild-type (15, 19, 20) (Fig. 4A). The presence of identical repeats in this region could promote amyloid formation by prioritizing intra-molecular stacking. To test this hypothesis, we expressed and purified a 12mer fragment termed 12merMUT, which contains 9 identical repeats, *i.e.*, 3 times more than 12mer[260:] (Fig. 4A and S5A,B). Strikingly, 12merMUT displayed extremely fast aggregation kinetics, orders of magnitude faster than 12mer[260:], even at >10-fold lower protein concentration (Fig. 4D and S5C). We could detect aggregate formation already during the purification (Fig. S5A,B). Unlike for the 1mer peptides or 12mer[260:], aggregation kinetics of 12merMUT were similar over a range of concentrations. Furthermore, the ThT fluorescence signal displayed a high scatter, suggesting that rapid polymerization could be leading to formation of large particles, as confirmed by fluorescent microscopy (Fig. 4E). By contrast, we were not able to detect any fibrils by EM or by AFM. This was also the case for 1mer[260:] or 1mer[656:]. Rapid polymerization of these fragments may lead to anisotropic growth and formation of large clumps, which could even be observed by eye in the test tubes.

To circumvent the problem of rapid aggregation and obtain fibrils comprised of identical repeats, we prepared constructs where 2, 3 or 4 copies of 1mer[260:] were fused to maltose-binding protein (MBP), followed by a tobacco etch virus (TEV) protease restriction site (Fig. 4F). The MBP tag increased the solubility of the tandem 33-aa identical repeats, and the fusion proteins were purified by affinity chromatography and tested in the ThT aggregation assay in the presence or absence of TEV to cleave off the N-ter tag. As expected, at about the same mass concentration of each MBP construct, that is the same concentration of 33-aa repeat, aggregation efficiency in the presence of TEV increased with the number of identical repeats in the construct, *i.e.*, from 2x to 4x (Fig. 4F). For the longest construct, MBP-4x1mer[260:], we observed aggregation by ThT fluorescence even in the absence of TEV (Fig. 4E, right panel). By negative stain EM, we could observe long fibrils in this sample (decorated with spherical buds likely representing uncleaved MBP), whereas in the +TEV sample, some shorter uniform fibrils were observed among more abundant and amorphous aggregates (Fig. 4G and S6A). No fibrils could be observed with the 2x or the 3x constructs. We obtained more abundant thick fibrils (∼20 nm in diameter) when MBP-4x1mer[260:] was purified using a longer induction to augment protein concentration, and treatment of these fibrils with pronase to shave off MBP reduced their thickness (5-8 nm in diameter) (Fig. S6B). Analysis of the MBP-4x1mer[260:] fibrils by cryo-EM yielded a similar classification as for the 12mer[260:] fibrils, with thick and thin polymorphs (Fig. S6C), suggesting that these fibrils contain a similar functional folding unit. We conclude that multiplication of identical repeats within the same peptide promotes fibril formation, which can be slowed-down by a globular tag.

Finally, we used budding yeast as a model to compare the behavior of 12mer[260:] and 12merMUT in a cellular environment. The budding yeast has been a useful model for studying the interaction of perilipins and the PLIN4 repetitive region with LDs (18, 30, 43, 44), and for studying the aggregation of human proteins such as α-synuclein or tau (33, 45, 46). Expression of 12mer[260:] and 12merMUT, tagged with mScarlet, from a strong constitutive promotor, did not affect the growth of wild-type yeast (Fig. S6D). Both constructs localized to LDs as well as to the plasma membrane after prolonged growth to accumulate LDs, as previously observed with 12mer[524:] (18), although 12merMUT accumulated on LDs somewhat more slowly than 12mer[260:] (Fig. S6E). Importantly, we could detect some large puncta of 12merMUT in a fraction of yeast cells (Fig. 4H). These puncta colocalized with the heat shock protein Hsp104, a protein disaggregase commonly used as a marker of protein aggregates (45, 47), suggesting that these structures represent aggregates of 12merMUT. Such puncta were never observed in yeast expressing the WT PLIN4 fragment 12mer[260:] (Fig. 4H, right panel). These results indicate that the identical repeats that are present in 9 copies in 12merMUT, compared to 3 copies in 12mer[260:], promote aggregation in a cellular context. We propose that the multiplication of identical repeats in mutant PLIN4 leads to more pronounced aggregation *via* intramolecular interactions that promote the growth of PLIN4 fibrils (Fig. 4I).

### Influence of lipid droplets on PLIN4 fibril formation

The PLIN4 repetitive region binds to LDs *in vitro* (Fig. 5A) (18, 28). We therefore wanted to assess how the presence of LDs affects PLIN4 fibril formation. We prepared artificial LDs composed of triglycerides, which form the neutral lipid core, and phosphatidylcholine (PC), the most abundant phospholipid forming the LD surface monolayer (48). In order to prepare a homogenous suspension of artificial LDs, we adjusted their density to be close to the density of water solution by the addition of brominated oil (28). We included in the LD preparation a small amount of fluorescently-labelled phospholipid (rhodamine-conjugated phosphatidylethanolamine; rhodamine-PE) to visualize the LDs by fluorescent microscopy. Incubation of these dense LDs with fluorescently-labelled PLIN4-12mer[260:] or 12merMUT showed that both proteins could bind to the LD surface, especially after a prolonged incubation (Fig. 5B).

**Fig. 5.**
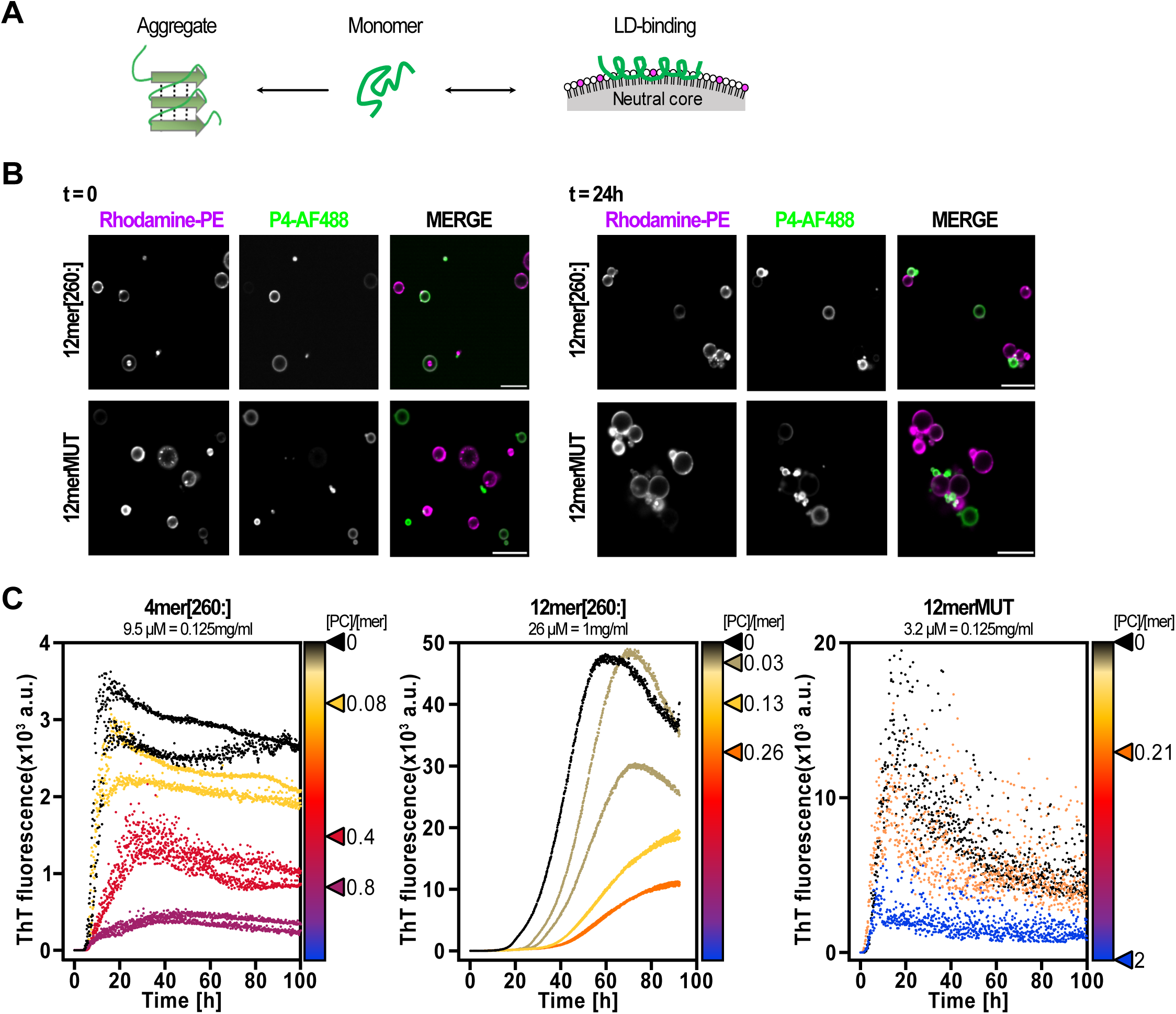
The influence of LDs on PLIN4 aggregation in vitro. (A) Schematic model of the different states of the PLIN4 repetitive region. The same region can interact with LDs or, more weakly, with lipid bilayers, or form amyloid fibrils. (B) Binding of 12mer[260:] or 12merMUT, labelled with AlexFluor488, to artificial LDs containing Rhodamine-PE was observed by confocal microscopy immediately or after a 24-hour incubation at RT. Scale bar: 10 µm. (C) ThT aggregation assay of 4mer[260:] (0.125 mg/mL), 12mer[260:] (1 mg/mL) and 12merMUT (0.125 mg/mL) in the presence of artificial LDs at the indicated lipid to protein molar ratio, expressed as the ratio of PC in the LD surface monolayer to 33-aa repeat for each construct. The exact ratios are presented with a color scale and indicated on the right side of each graph. Protein concentrations were chosen such as to obtain good aggregation in the absence of LDs. See also Fig. S7 for comparison of aggregation in the presence of liposomes.

Next, we performed a ThT aggregation assay with 4mer[260:], 12mer[260:] or 12merMUT in the absence or presence of an increasing concentration of LDs. We calculated the LD to protein ratio in the reactions by using the molar PC concentration as a measure of available LD surface, which we then normalized to the total molar concentration of 33-aa repeats (mer) in each reaction (Fig. 5C). The aggregation of 4mer[260:] or 12mer[260:] was slowed-down or completely blocked, respectively, by an increasing concentration of LDs (Fig. 5C). By contrast, the aggregation of 12merMUT was much less affected by LDs at the same PC (surface lipid) to protein molar ratio (Fig. 5C, right-most panel). Furthermore, PC liposomes (bilayer membranes) had less effect on aggregation of 4mer[260:] and 12merMUT compared to LDs, even at a >100-fold higher PC concentration (Fig. S7). This result correlates with a lower affinity of PLIN4 amphipathic helix for bilayers compared to LDs (18, 28). We therefore conclude that aggregation of the PLIN4 repetitive region may be slowed down by LDs via a competitive mechanism: for WT PLIN4 repetitive region, binding to LDs is more pronounced than aggregation, whereas in the case of PLIN4-12merMUT, aggregation is less sensitive to the presence of LDs, likely because aggregation occurs faster than LD binding.

## Discussion

Many degenerative diseases are linked to protein misfolding and amyloid deposits in affected tissues. The identity of the protein and the 3D structure of the protein deposit are intimately linked to each specific disease (4, 49). Here, we show that the LD-binding protein PLIN4 is a novel amyloid-forming amyloid, as demonstrated by cryo-EM of the purified PLIN4 repetitive region. We further show that aggregation is strongly promoted by multiplication of identical 33-aa repeats found in the MRUPAV vacuolar myopathy patients (15), whereas the multiplication does not affect the interaction of PLIN4 with LDs in cells. The genetically-encoded expansion of PLIN4 may therefore represent a toxic a gain-of-function mutation that promotes aggregation of PLIN4 in skeletal muscle, where it is highly expressed. This is consistent with the observations of PLIN4 accumulation in skeletal muscle tissue of patients carrying the *PLIN4* expansion, and of colocalization of PLIN4 with p62 and other aggrephagy markers (14, 15, 19–21). Our biochemical analyses suggest that the enhanced aggregation propensity of mutant PLIN4 arises primarily from repeat identity rather than from the increase in repeat number *per se*, raising broader implications for how repetitive protein architectures can influence amyloid assembly.

The kinetic assays suggest that the disease expansion in PLIN4 strongly promotes fibril formation via intramolecular interactions between identical repeats. This model is in agreement with the large increase in aggregation kinetics when comparing two PLIN4 fragments of the same length (12 x 33 aa = 396 aa) but differing by the number of identical repeats, as well as the loss of concentration dependence with the increase in identical repeats (Fig. 4D). Furthermore, when we multiplied the number of identical repeats within the same peptide (from 2 to 4), the aggregation kinetics was increased even though the total number of identical repeats was kept similar between the reactions (Fig. 4F). One possibility is intramolecular stacking, which has been observed for the fungal protein Het-S, whereby two imperfect repeats of 21 aa within the same molecule form a cross-β stack (50, 51), and for the functional amyloids curli secreted by many Gram-negative bacteria, including CsgA of *E. coli,* where intra-molecular stacks are composed of multiple imperfect repeats (52, 53). Alternatively, the large clumps observed with the 12merMUT construct suggest that identical repeats could promote anisotropic growth of the amyloid structures. However, our results also indicate that the presence of identical repeats is not required for formation of PLIN4 amyloids. A number of individual PLIN4 repeats can assemble into filaments, whereas the variable residues within some repeats could represent anti-aggregation residues. The presence of “gate-keeper” residues within a few central repeats in CsgA reduces amyloid formation (54, 55); similarly, the presence of specific flanking sequences can inhibit the aggregation of polyglutamine repeats (56). The presence of identical repeats in PLIN4 mutant may therefore abolish the anti-aggregation properties of flanking repeats. These considerations also suggest strategies for inhibiting PLIN4 aggregation by the use of interfering peptide (57, 58), and will be the subject of future studies.

Another intriguing question is the relationship between PLIN4 binding to LDs and amyloid formation. While the PLIN4 repeat expansion does not significantly impact its interaction with LDs in cellular models, we show that LDs inhibit PLIN4 aggregation *in vitro*, suggesting that they could act as a buffer to decrease the propensity of PLIN4 to aggregate. Further studies on the relationship between PLIN4 binding to LDs and its aggregation may offer other therapeutic strategies for MRUPAV patients. In this respect, it is also important to ask why the PLIN4 expansion appears to specifically affect skeletal muscle tissue, whereas no defect in the function of adipose tissue, where PLIN4 is also highly expressed, has been observed. In adipocytes, PLIN4 is well adapted to handling fast forming LDs (28). In muscle, LDs are abundant within myofibers and play an important role in muscle contraction, with LD number, distribution, and association with mitochondria, being highly variable (24, 25). Muscles are prone to rapid changes in lipid composition owing to diet-induced changes (24, 59). Differences in lipid metabolism may favor or disfavor PLIN4 aggregation versus its normal function on LDs. Muscle cell and animal models will be needed to address these questions.

According to currently available data, MRUPAV is an extremely rare genetic disease, with only a few families reported world-wide (15, 19, 20). However, the genetic expansion of highly similar repeats in *PLIN4* cannot be identified by the high-throughput screening approaches routinely used in diagnostic, and recent studies suggest that toxic expansions in *PLIN4* may be considerably more common (21, 60), underlying the importance of understanding the molecular mechanism of PLIN4 aggregation. There are other diseases associated with protein aggregation within muscle fibrils, in particular myofibrillar myopathies, which often include aggregates of the muscle-specific intermediate filament desmin, which are a likely to be amyloid in nature (13, 61). Furthermore, degenerative diseases are often linked to defects in cellular lipid metabolism and changes in membrane lipid composition during ageing (62, 63). LDs have been suggested to act as a buffer for toxic protein aggregates and were shown to suppress lipotoxic effects associated with Parkinson’s disease and other degenerative disorders (34). In cells of Parkinson’s and Alzheimer’s models, LDs and other lipid-bound organelles have been observed in proximity of amyloids (33, 35). Different lipids promote or inhibit aggregation or amyloid formation of amyloid-β or α-synuclein, and lipids can be incorporated into amyloid fibrils (64–66). By being expressed in two tissues with different lipid metabolic traits, PLIN4 provides a favorable context to connect *in vitro* studies performed using pure proteins and lipids to cellular investigations, clinical observations, and model organisms for pathology.

Our current study is clearly limited by being largely based on biochemical observations using fragments of PLIN4. The size and disordered and repetitive nature of PLIN4 represent significant challenges towards studying the behavior of the full-length protein *in vitro,* as well as towards establishing a muscle-based cellular model to study PLIN4 aggregation. These will be the goals of our future studies.

## Acknowledgments

We thank Manuel Giménez Andrès for assistance with protein purification and labelling, Amel Balhoul for help with CD spectroscopy, Laura Picas for advice on the AFM experiments, Véronique Albanèse for advice and plasmids for yeast experiments, and Bruno Antonny, Andrey Kajava, Katarina Trajković and Dimitris Xirodimas for helpful discussions and comments on the manuscript. This work benefited from the CryoEM platform of I2BC, supported by the French Infrastructure for Integrated Structural Biology (FRISBI) [ANR-10-INSB-05-05], the MRI imaging facility, a member of the national infrastructure France-BioImaging, supported by the French National Research Agency (ANR-10-INBS-04, Investissements d’avenir), the joint IGMM-CRBM “yeast media and technologies service”, and the Synbio3 platform (IBMM, Montpellier University, France), supported by GIS IBISA. The NanoImaging Core at Institut Pasteur is acknowledged for support with sample preparation, image acquisition and analysis. The NanoImaging Core was created with the help of a grant from the French Government’s Investissements d’Avenir program (EQUIPEX CACSICE - Centre d’analyse de systèmes complexes dans les environnements complexes, ANR-11-EQPX-0008). This work was supported by the European Research Council (ERC Synergy 856404, SPHERES), the Agence Nationale de la Recherche (ANR-23-CE44-0026) and by AFM Téléthon (SR 2024 #289888).

## Author contributions

Conceptualization: A.C., A.R. and L.B.; Data curation: N.F., C.M., C.F., S.L., L.B., A.C.; Formal analysis: C.C., S.L. and L.B.; Funding acquisition: A.C., A.R. and L.B.; Investigation: C.C., N.F., B.S., C.M., C.F., S.L., T.P., S.P., L.B., A.C.; Methodology: N.F., C.M., S.L., L.B., A.R., A.C.: Project administration: A.C., Resources: C.M., A.R, and S.R., Supervision: A.C., L.B., S.L.; Visualization: C.C., N.F., B.S., C.M., C.F., S.L., L.B.; Writing – original draft: A.C.; Writing – review & editing: L.B., C.C., N.F. C.M., and A.R.

## Declaratiton of interests

The authors declare no competing interests.

## Supplemental information

Document S1. Table S1, Figures S1-S7.

Table S2. Excel file containing descriptions, amino acid and DNA sequences for all proteins and peptides used in this study, related to Key Resources Table.

Table S3. Excel file containing DNA and transcribed amino acid sequences of synthetic genes used for construction of mammalian PLIN4 expression plasmids.

## Methods

### Protein sequence analysis

The Uniprot entry Q96Q06 was used as the default sequence for human PLIN4. The aa repeats were identified using HHrepID rom the MPI Bioinformatics Toolkit as previously described (30, 67). Repeats were aligned using AlignmentViewer (http://alignmentviewer.org) and the repeat alignment was presented using Weblogo (68). The TAPASS pipeline (40), including amyloid prediction algorithms ArchCandy2, Pasta and Tango (Ahmed et al., 2015; Fernandez-Escamilla et al., 2004; Walsh et al., 2014), was used to identify the ARs within the PLIN4 sequence. The amyloidogenic score of individual 33-aa repeats was determined using ArchCandy2.

### Plasmids for mammalian cell culture

Plasmids for stable expression of full length wild-type PLIN4 (pSB-PLIN4-WT-GFP, pSR002) or PLIN4 mutant (pSB-PLIN4-MUT-GFP, pSR003) in HeLa cells were constructed using the Sleeping Beauty (SB) transposon system (69). Four synthetic genes (GeneArt Gene Synthesis, Thermo Fisher Scientific), listed in Table S3 (pSG-Nter, pSG-RPT1, pSG-RPT2, pSG-Cter-GFP), encoding codon-optimized human PLIN4 and encompassing the entire wild-type sequence with a C-terminal GFP tag, were assembled using the NEBuilder DNA HiFi assembly kit (New England Biolabs). The resulting construct was cloned into the pSBtet-Pur vector (a gift from Eric Kowarz, Addgene plasmid#60507). The PLIN4 mutant construct was generated from the wild-type sequence by inserting the repeat expansion (pSG-12mer-id, see Table S3) using the same NEBuilder kit and cloned into the same vector. Plasmids for transient expression of PLIN4 constructs with deletions in the repetitive region were constructed by amplifying the regions of interest from the same synthetic genes and cloned with NEBuilder kit into the peGFP-N1 plasmid (Addgene plasmid #6085-1). Plasmids containing 19 repeats were obtained fortuitously during cloning of the full-length plasmid. Plasmids sequences were verified by whole-plasmid Nanopore sequencing.

### Establishment of inducible stable cell lines and transiently transfected cells

HeLa cells (ATCC# CCL-2), Huh-7 cells (provided by M. Moriel-Carretero, CRBM, Montpellier) and Hep3B cells (provided by P. Corbeau, IGH, Montpellier) were grown in Dulbecco’s modified Eagle’s medium (DMEM) supplemented with 4.5 g l−1 glucose (Life technologies), 10% fetal bovine serum (FBS, Life technology) and 1% Penicillin/Streptomycin antibiotics (Life technologies). Sub-confluent cells were co-transfected with plasmids encoding either pSB-PLIN4-WT-GFP (pSR002) or pSB-PLIN4-MUT-GFP (pSR003), along with the Sleeping Beauty transposase SB100X (pCMV(CAT)T7-SB100; a gift from Zsuzsanna Izsvak, Addgene plasmid # 34879) (70) using Fugene HD in Optimem medium (Life technologies). Twenty-four hours post-transfection, stable integrants were selected in tetracycline-depleted medium supplemented with puromycin (1 µg/mL). Inducible expression of the transgenes was triggered by adding doxycycline (1 µg/mL) overnight and confirmed *via* Western blot analysis and immunofluorescence microscopy. For observation by confocal fluorescent microscopy, stably transfected cells were induced with doxycycline (1 µg/mL) overnight in standard growth medium or standard growth medium containing 150 μM oleic acid (Sigma) in complex with fatty-acid free BSA (Sigma). LDs were stained with Lipi-Blue (0.1 µmol/L, Dojindo) for 30 min in the standard growth medium without phenol red immediately prior to live imaging by confocal microscopy. For transient expression of PLIN4-GFP constructs, sub-confluent HeLa cells were transfected with peGFP-N1 derivated plasmids expressing PLIN4 variants fused to GFP (pCM144, pcM144, pCM152, pCM162) using Fugene HD in Optimem medium (Life technologies). 24h post transfection, LDs were stained with Lipi-Blue (0.1 µmol/L, Dojindo) for 30 min in standard DMEM without phenol red prior to live cell imaging on a spinning disc microscope. For fluorescence recovery after photobleaching (FRAP) experiments, cells were treated as for transient transfection.

### Analysis of PLIN4 expression and localization by Western blot and immunofluorescence (IF)

For Western blot analysis, stably transfected cells were induced with doxycycline (1 µg/mL) overnight in standard growth medium or standard growth medium containing 150 μM oleic acid (Sigma) in complex with fatty-acid free BSA (Sigma). Whole-cell protein extracts were prepared in RIPA buffer (50 mM Tris⋅HCl, pH 7.5, 150 mM NaCl, 1% NP-40, 0.5% sodium deoxycholate, and 0.1% SDS) supplemented with protease inhibitors (Complete, Roche) and phosphatase inhibitors (PhosSTOP, Roche). Protein concentrations were determined using a Bradford assay. Equal amounts of protein were resolved by SDS–PAGE, transferred to PVDF membranes, and incubated with the indicated primary antibodies: anti-GFP (Roche; 11814460001; 1:1,000) and anti–α-tubulin (clone DM1A; Sigma; T6199; 1:5,000). Membranes were then incubated with HRP-conjugated anti-mouse IgG secondary antibodies (Thermo Fisher Scientific), and signals were detected using enhanced chemiluminescence (Luminata Crescendo, Merck Millipore) with Odyssey imaging system (Odyssey M, LI-COR). For IF analysis, cells were fixed in 3.2% paraformaldehyde (Thermo-28906) for 15 min at room temperature (RT). They were then gently permeabilized with 0.5% saponin in phosphate-buffered saline (PBS) containing 0.5% FBS for 15 min at RT, washed with PBS and incubated in a blocking solution containing PBS and 0.5% FBS for 30 min at RT. They were then incubated with primary antibody cocktails using antibodies against TOMM20 (Abcam-ab56783, mouse monoclonal, diluted 1/250) and against PLIN4 (Abcam-ab234752, rabbit polyclonal, diluted 1/250) overnight at 4 °C, rinsed in PBS and incubated with secondary antibodies and Lipi-Blue stain (0.1 µmol/L, Dojindo) for 1 hour at RT, protected from light, and mounted for imaging.

### Cloning and plasmid construction for protein purification

All purified PLIN4-derived proteins were expressed from pET21b (Novagen)-based plasmids constructed using synthetic genes encoding human sequence (Uniprot entry Q96Q06), codon-optimized for human PLIN4 and to reduce the percent of sequence identity, ordered from Eurofins (designed using the Eurofins algorithm). All PLIN4-4mer and PLIN4-12mer sequences were subcloned into pET21b using NheI and HindIII restriction sites, to be expressed without an N-ter or a C-ter tag. The only exception was 4mer[260:], which was inserted into pET21b digested with BamH & HindIII, preserving the N-ter T7 tag of 13 aa (1.4 kDa). PLIN4-20mer was cloned into pET21b as previously described (18), for expression and purification without a tag. PLIN4-1mer synthetic peptides were ordered from Proteogenix. To produce the MBP- 1mer[260:] fusion proteins, the indicated number of 1mer[260:] repeats (2, 3 or 4) was amplified by PCR from plasmid pCC05, and inserted into the pETM40 bacterial expression vector using NcoI/StuI restriction site to produce in-frame fusion with a 5’ MBP tag-encoding sequence, followed by the TEV protease cleavage site (5’-GAGAATCTTTATTTTCAGGGC-3’). All proteins and peptides used in this study are described in detail in Table S2, and the names of plasmids used for their purification are provided.

### Yeast plasmids, growth and media

To construct plasmids for expression of 12mer[260:] (pCM86) and 12merMUT (pCM85) in budding yeast, sequences indicated in Table S2 (codon-optimized for humans) were subcloned by Gibson cloning into a pRS416 (*CEN, URA3*, Amp^R^) plasmid containing the strong GPD promoter (71). They were then tagged at the C-termini with an mScarlet fluorescent protein (DNA amplified from pBBK83, addgene #179067) using Gibson cloning (Linker sequence: GSGPSGPTG). The yeast strain used for all experiments was BY4741 MATa his3Δ1 leu2Δ0 met15Δ0 ura3Δ0 *HSP104::GFP::HygrMX*. The GFP-coding cassette (72) was inserted at the 3’ end of *HSP104* by homologous recombination and verified by fluorescent microscopy. Yeast cells were grown in yeast extract/peptone/glucose (YPD) rich medium, or in synthetic complete (SC) medium containing 2% glucose. Yeast was transformed by standard lithium acetate/polyethylene glycol method. For observation of aggregates, liquid cell cultures were inoculated from a single colony and grown for 18 h at 30 °C in SC-Ura with 2% glucose to maintain plasmid selection and observed directly by confocal microscopy. For observation of LDs, yeast cultures were grown under the same growth conditions for 24h or 48h. LDs were stained with AutoDOT (CliniScience) for 20 min at room temp in growth medium and the cells were washed twice before observation.

### Protein expression and purification

All proteins and peptides used in this study are listed in Table S2. The recombinant PLIN4-20mer, PLIN4-12mer[260:], PLIN4-12mer[524:], PLIN4-12merMUT, PLIN4-4mer[260:], PLIN4-4mer[326:], PLIN4-4mer[557:] and PLIN4-4mer[689:] proteins were purified without a tag following a previously-established procedure (18). They were expressed in *E. coli* BL21(DE3) at 37°C for 1 h 40 min after induction with 1mM IPTG (at OD_600_ = 0.6-0.8). Always at 4°C, cells were lysed in lysis buffer A (20 mM Tris-HCl pH 7.5, 10mM NaCl, 1mM dithiothreitol (DTT), 5mM PMSF, protease inhibitors tablet cOmplete; Roche) by sonication (amplification 40%, 10 sec ON, 20 sec OFF 10 times) and the lysates were centrifuged at 40,000 g for 30 min. Boiling step of the supernatants at 95°C was performed for 30 min in a water bath, followed by 10 min in ice and centrifugation at 40,000 g for 20 min. 2 rounds of dialysis (90 min each) were performed against lysis buffer B (20 mM Tris-HCl pH 7.5, 10 mM NaCl, 1 mM TCEP) using Spectra/Por membranes with a cut off of 6000 Da (Spectrum labs), followed by 30 minutes of centrifugation at 40,000 g. Purification was performed on a NGC FPLC system (BioRad), using a Hiprep SP 16/10 cation-exchange column (Cytiva). Proteins were eluted with a linear salt gradient with elution buffer B (20 mM Tris-HCl pH 7.5, 1M NaCl, 1mM TCEP) and protein-containing fractions were pooled, aliquoted for single use and stored at -80°C. Protein concentration was determined by densitometry from SDS-PAGE gels stained with GelCodeBlue (Thermofisher) using bovine serum albumin (BSA) standards for calibration. Note that due to the lack of aromatic residues in the PLIN4 repetitive region, protein concentration cannot be determined by standard assays such as Bradford; for details see (18).

MBP-fusion proteins containing PLIN4-1mer[260:] repeats (2, 3 or 4, as indicated) were expressed in *E. Coli* Arctic (DE3) cells at 18°C overnight following induction with 1mM IPTG (at OD_600_ = 0.6-0.8). Cells were lysed by sonication in lysis buffer A (20 mM Tris-HCl pH7.5, 50 mM NaCl), supplemented with 10% glycerol, 1 mM TCEP, 1 mM EDTA, 0.5 mM PMSF, 1.5 μM pestatin, 2 μM bestatin and a protease inhibitors coctail cOmplete (Roche), as described above, and centrifuged to produce cleared lysates. The lysates were incubated for 2 h at 4°C under rotation with amylose beads and then sequentially eluted with 1 mL of maltose elution buffer (lysis buffer supplemented with 50 mM maltose). Elution fractions 1 and 2 were pooled together and immediately used in the ThT aggregation assay (immediate use without freeze-thawing was essential to reduce protein degradation). Protein concentration in the eluted fractions was determined by densitometry from SDS-PAGE gels stained with GelCodeBlue using bovine serum albumin (BSA) standards applied on the same gel for calibration. Similar protein concentrations were obtained with the different MBP-constructs, as indicated in the figure legends. TEV (Tobaco Etch Virus) protease was purified in-house and added to the ThT assays at the same time as the MBP-fusion proteins at a concentration of 0.05 mg/mL, when indicated.

### Protein fluorescent labelling

Purified PLIN4 fragments were labelled via endogenous cysteines using Alexa Fluor 488 C5 maleimide dyes (ThermoFisher), as previously described (18). Briefly, for proteins purified in the presence of DTT, they were first dialyzed on a NAP10 desalting column (Cytiva) against 50 mM Tris-HCl pH 7.5, 120 mM NaCl buffer The reactions were carried out with 5-fold molar excess (per Cys) of the dye for 20 min at 4°C, then blocked with 2mM DTT. The free dye was removed using desalting NAP-5 columns (Cytiva) equilibrated with 50 mM Tris-HCl pH 7.5, 120 mM NaCl buffer, supplemented with 2 mM DTT. The fractions were analyzed by SDS-PAGE and UV-visible chromatography. Labeled proteins were stored at -80°C in small aliquots and mixed with a 5-fold excess of identical unlabeled protein when needed.

### Synthetic peptides

Unmodified synthetic peptides of 33 aa were ordered from Proteogenix. They were stored in 1 mg aliquots at -20°C and dissolved in dH_2_O or in 25 mM Tris, pH 7.5, 100 mM NaCl just before use.

### ThT aggregation assay

Immediately prior to an aggregation assay, in order to remove any aggregation seeds (oligomers or pre-formed fibrils), all proteins and synthetic peptides were centrifuged at 100,000 g for 1 h at 4°C. Aggregation was performed in 25 mM Tris, pH 7.5, 100 mM NaCl and 1 mM DTT (freshly added), supplemented with 15 µM Thioflavin T. The reaction mixtures (250 μl) were pipetted into wells of a Greiner BlackPlate 96 Flat containing one 2 mm glass bead per well and sealed with a plastic seal. Fluorescence was monitored continuously every 10 min in a TECAN SPARK fluorescent plate reader at 37°C, with orbital shaking for 300 s, amplitude of 2.5 mm and frequency of 216 rpm. ThT fluorescence was acquired in the top reading mode at 2x2 multiple reads per well, with a 430/485 nm filter, 40 % manual gain, 50 % mirror, 30 flashes and 40 µs of integration flash, at an optimized Z-Position (usually around 18500 µm). After aggregation curves reached a plateau, samples were removed from the plate and stored in low-binding Eppendorf tubes in the dark at room temperature.

### Protein pelleting assay

When the ThT fluorescence signal of aggregation assays with different PLIN4 constructs reached the plateau, 50 µL of each reaction was centrifuged at 40 K rpm for 30 min at 25°C. Supernatants were collected and mixed with 2x Laemmli buffer containing 5 mM DTT, and pellets were resuspended in 100 µl of the 1x Laemmli buffer with DTT. Equal volume (10 µl) of each sample was loaded on a 17% SDS polyacrylamide gel and analyzed by SDS polyagrylamide gel electrophoresis (SDS PAGE) in Tris-glycine migration buffer containing 0.05% SDS. Proteins were stained using GelCode Blue (Thermo Scientific).

### Circular Dichroism

The experiments were conducted on a Jasco J-815 spectrometer at room temperature with a quartz cell of 0.05 cm path length. Each spectrum is the average of several scans recorded from 200 to 260 nm, with a bandwidth of 1 nm, a step size of 0.5 nm, and a scan speed of 50 nm/min. Control spectra of 50 mM Tris, pH 7.5, and 150 mM NaCl buffer without or with 50% TFE were subtracted from the protein spectra.

### Negative stain EM

Negative stain EM of amyloid fibrils was performed as described (73). Briefly, PLIN4 samples were diluted to 0.1 mg/mL in MilliQ water. Glow discharged carbon coated 400 mesh copper grids were floated over 30 µL drop for 1 min. The excess of liquid was absorbed on Whatman filter paper, and the grid was stained for 1 min over a 30 µL droplet of 1% uranyl acetate. The grids were observed under Jeol 1400 electron microscope at 80 kV and 10 K magnification, images were recorded on a Gatan Rios CCD camera with automatic drift correction.

### Cryo-EM

PLIN4-12mer[246] filament preparations were screened by negative stain EM to identify the most homogenous samples suitable for Cryo-EM. Quantifoil R3.5/1 grids were prepared using 3µl of 1mg/ml plin4 sample using a Vitrobot. Data collection on Titan KRIOS equipped with Falcon 4i camera at the Institut Pasteur. 6000 images were acquired with a 0.77 Å pixel size and defocus ranging from 0.4 to 2.5 µm. All data were processed with Relion4 (74) for 3D classification using a 1024x1024 pixel box size binned to 256, or an unbinned 350x350 pixel box size to detect the 4.7 Å spacing.

### Atomic force microscopy

AFM experiments were done using a JPK NanoWizard IV xp microscope (Bruker nano GmbH, Germany) using the PeakForce tapping mode in liquid and PeakForce-HIRS-F-B probes (nominal tip radius = 1 nm, nominal spring constant 0.1 N/m, Bruker). PLIN4-12m[260:] fibrils [1 - 2 mg/mL], collected after Thioflavin T assay, were diluted ten times then incubated for 10 min on freshly cleaved mica surface before imaging in buffer (25 mM Tris, pH 7.5, 100 mM NaCl). 256 x 256 pixels images were collected at scan rate of 5Hz, with a PeakForce setpoint of 300pN and Z-Range of 7.5 µm. JPK data analysis software (version 8.0.177) was used to process the image data by flattening the height topology and to measure manually the periodicity from the height profile.

### Lipids

DOPC (1,2-dioleoyl-sn-glycero-3-phosphocholine), diphytanoyl-PC (1,2-diphytanoyl-sn-glycero-3-phosphocholine), diphytanoyl-PS (1,2-diphytanoyl-sn-glycero-3-phospho-L-serine) and Rhodamine-PE (L-α-Phosphatidylethanolamine-N-(lissamine rhodamine B sulfonyl) were purchased from Avanti Polar Lipids as chloroform solutions. Triolein (1,2,3-tri-(9Z-octadecenoyl)-glycerol) and BVO (Brominated Vegetal Oil, CAS: 8016-94-2) were purchased from Sigma-Aldrich and Spectrum Chemical MFG Corp, respectively. BVO was purified before use as described (28). Lipids were stored as chloroform stock solutions at -20°C under argon.

### Liposome preparation

Phospholipids in chloroform were mixed at the desired molar ratio and dried under argon flux. To prepare fluorescently-labeled liposomes, 0.1 % of Rhodamine-PE was included in the mixture. The lipid film was hydrated in 1 mL of HK buffer (50 mM HEPES, pH 7.4, and 120 mM K-Acetate) to a concentration of 2 mM phospholipid and freeze-thawed five times in liquid nitrogen. Prior to use, 500 µL of liposomes were extruded 19x through a polycarbonate filter of 0.2 µm pore size using a mini-extruder. Extruded liposomes were stored at room temp and used within 2 days following extrusion.

### Limited proteolysis

Prior to the experiment, proteins were centrifuged 10 min at 20,000 g, 4°C, to remove any precipitate, and diluted in HK buffer (50 mM HEPES, pH 7.4 and 120 mM K-Acetate) containing 1 mM MgCl_2_ to 0.1 mg/mL was mixed or not with diphytanoyl (50% PC, 50% PS) liposomes at 1:1000 protein-to-phospholipid molar ratio in 120 µL final volume. The mix was incubated for 10 min at 25°C with 300 rpm agitation prior to addition of 0.2 µg/mL of subtilisin. At indicated times, 20 μL of the reaction was removed and the reaction was stopped immediately with 2mM PMSF. Samples were analyzed by SDS-PAGE and proteins were stained with Silver stain.

Limited proteolysis of fibrils formed by MBP-PLIN4 constructs (1mg/ml) was carried out using pronase at final concentration of 0.1 mg/ml for 30 minutes at 20°C in the assembly buffer. Pronase digestion was confirmed by SDS-PAGE where the MBP-PLIN4 fusion protein band completely disappear within 30 minutes. At 30 minutes an aliquot was withdrawn immediately to prepare electron microscopy grids for negative stain and cryo-EM, as described.

### Artificial LD preparation

Artificial LDs with 13.3 µM DOPC were prepared in 50 mM HEPES, pH 7.4, and 120 mM K-Acetate (HK) buffer and extruded through polycarbonate filters of 8-µm pore size using a mini-extruder (Avanti Polar Lipids), as described (28). All lipids were from Avanti polar lipids, except TG(18:1/18:1/18:1) and BVO (CAS: 8016-94-2), which were from Sigma and Spectrum Chemical MFG Corp, respectively. Contaminants such as free FA were removed from BVO as described (28). Purified BVO was dissolved in chloroform and stored at -20°C under argon. To prepare aLDs of defined density, close to the density of the HK buffer, and diameter, we used to protocol described in (28). The aLD density was verified experimentally by confocal microscopy. For a volume fraction of 0.75% oil in buffer, obtaining a suspension of aLDs with a calculated diameter of 10 µm, adapted for visual analysis by confocal fluorescent microscopy, requires a concentration of phospholipid [PL] = 12.5 µM. To prepare the aLDs, we first mixed 10 µL of triolein (9.11 mg) and 5 µL of BVO (6.6 mg) from stock solutions in chloroform (≈ 90 mg/ml). For 10 µm aLDs, the mix was supplemented with 25 nmol of DOPC (18:1/18:1)) and 0.125 nmol Rhodamine-PE for microscopy experiments. After evaporation of chloroform under a stream of argon (∼30 min using a low flow rate), the final oil droplet containing the phospholipids was resuspended in 2 mL HKMD buffer (HEPES 50 mM pH 7.2, K acetate 120 mM, MgCl2 1 mM, DTT 1 mM), hence leading to a 0.75% oil suspension. The suspension was briefly vortexed and pipetted 5x with a Hamilton syringe, then extruded 19 times through 8 µm polycarbonate filters using a hand mini extruder (Avanti). Note that extrusion through this pore size yields aLDs approximately 10 µm in diameter when assessed by dynamic light scattering (28). The aLDs were kept at room temperature under argon, protected from light, and used on the same day.

### Analysis of protein binding to artificial LDs

In a clean glass tube, 500 µL final volume of freshly prepared artificial LD suspension was gently mixed with 0.25 µM of protein, where 1/5^th^ is fluorescently labelled with AlexaFluor-488, in a final volume of 600 µl (in HK buffer supplemented with 1mM MgCl2 and 1mM DTT). The binding of proteins to LDs was analyzed by confocal fluorescent microscopy immediately or after a 24h incubation at room temp. 30 µL of the LD-protein mixture was gently deposited on an 8-well Ibidi slide and containing 80 µL of buffer, incubated for 15 min to allow the LDs to attach to the surface of the slide and observed by a spinning disk microscope.

### ThT aggregation assay in the presence of LDs or liposomes

Before experiment, aliquot of unilamellar DOPC liposome suspension (2 mM DOPC) were thawed at room temperature and then extruded through a 0.4 µm diameter filter. For LDs, 13.3 µM DOPC aLDs were freshly prepared and extruded through an 8 µm diameter filter. Different volumes of aLDs (0 µL, 10 µL, 50 µL or 100 µL) or liposomes (0 µL, 10 µL or 100 µL) were added to wells of a Greiner BlackPlate 96 Flat along with different PLIN4 fragments, as indicated: 4mer[260:] (0.125 mg/mL, [mer] = 9.5 µM), 12mer[260:] (1 mg/mL, [mer] = 26 µM) or 12merMUT (0.125mg/mL [mer] = 3.2 µM). The protein concentrations were chosen such as to assure good aggregation kinetics and minimize the total amount of protein required, based on previous experiments in the absence of lipids, and thus differed between different constructs. For each condition, the molar ratio between [PC] and individual 33-aa PLIN4 repeats [mer] was calculated. Note that the PC concentration of liposomes is >100x higher than in the case of LDs, due to the smaller size of liposomes. As the liposomes contain a lipid bilayer, compared to the LD monolayer, the PC concentration in the case of liposomes was divided by 2 to obtain the total PC area available for protein binding, assuming that the liposomes remain intact during the course of the experiment. This is difficult to verify due to adsorption of liposomes/LDs to the sides of the wells during the course of the experiment.

### Confocal microscopy and image analysis

Multi-dimensional images of PLIN4-GFP in live Huh7 and Hep3B cells were acquired at 37°C with an LSM980-NLO confocal microscope (Zeiss), using a 63X Plan-Apo 1.4NA oil-immersion objective. The microscope is equipped with T-PMT camera and driven by Zeiss Zen Blue software. Excitation sources used were: 405 nm diode laser and an Argon laser for 488 nm and 514 nm. The same set-up was used for imaging of fixed cells by IF. Images were analyzed using ImageJ and data was plotted using GraphPad (Systat Software).

Observations of PLIN4-GFP constructs in live HeLa cells were performed with an inverted Olympus Ixplore Spin SR microscope coupled with a spinning disk CSU-W1 head (Yokogawa) using 60X UPLXAPO 1.42 NA DT 0.15 mm oil-immersion objective. Z stacks of 10 images with a step size of 0.5 µm were acquired with sCMOS Fusion BT Hamamatsu camera, using 488 nm and 561 nm (100 mW) lasers with GFP Narrow filter. The system was driven by CellSens Dimension 3.2 software. FRAP was performed using a 488nm (100mW) FRAP laser. A circular area on a single LD or LD cluster was selected; three images were taken before bleaching, followed by a post-bleach time-course of 1 image/s (note that the rate of image acquisition was limited by the low fluorescent signal). Background fluorescence outside the cell was subtracted from the bleached area and the signal was normalized to the whole cell signal for each time-point. Data was processed with ImageJ and Excel and plotted using GraphPad.

Observations of artificial LDs and budding yeast were performed using the same spinning-disk set-up. Z stacks of 10 images with a step size of 0.5 µm were acquired with sCMOS Fusion BT Hamamatsu camera, using 488 nm and 561 nm (100 mW) lasers with GFP Narrow filter. Observations of PLIN4-12mer[260:] and PLIN4-12merMUT aggregates after ThT assay were performed with an Axioimager Z2 Upright Zeiss microscope using 40X EC Plan Neofluar 1.4 NA oil-immersion objective. Images were acquired with scMOS ZYLA 4.2 MP camera and a CFP filter. The system was driven by Metamorph software. Images were analyzed using Image-J. Aggregates in yeast were quantified manually. Data was plotted using GraphPad.

## Statistical analysis

All statistical analyses were performed using GraphPad Prism (ver. 10.6.1)softwre (GraphPad Software, Boston, Massachusetts, USA). For cross-analysis, a Shapiro test was used to control for normality of the samples, followed by a pair-wise t-test or a Mann-Whitney test. Details are described in the respective figure legends.

