## Supplemental Data: Table S1, Figures S1-S7 for "Perilipin 4 repetitive region forms amyloid fibrils promoted by a genetic expansion found in myopathy patients"

**Choufani, Fuggetta et al.,**

### Supplementary Information, Document S1

Table S1,
Figures S1-S7

Table S1. Analysis of individual 33-aa PLIN4 repeat sequences.

| First residue[a] | Repeat # | % identity to r6[b] | % indentity to cons. seq.[c] | ArchCandy score[d] | Repeat sequence[e] |
|---|---|---|---|---|---|
| 95 | r1 | 42.4 | 48.5 | 0.553 | VCSKMSR**AKDAVSSGVASVVDVAKGV**VQGGLDT |
| 128 | r2 | 60.6 | 66.7 | 0.557 | TR**SALTGTKEVVSSGVTGAMDMAKGAV**QGGLDT |
| 161 | r3 | 78.8 | 78.8 | 0.519 | SKA**VLTGTKDTVSTGLTGAVNVAKGTVQAG**VDT |
| 194 | r4 | 75.8 | 81.8 | 0.519 | TKT**VLTGTKDTVTTGVMGAVNLAKGTVQTG**VET |
| 227 | r5 | 75.8 | 72.7 | 0.486 | SKA**VLTGTKDAVSTGLTGAVNVARGSIQTG**VDT |
| 260 | r6 | 100 | 87.9 | 0.519 | SKT**VLTGTKDTVCSGVTGAMNVAKGTIQTG**VDT |
| 293 | r7 | 100 | 87.9 | 0.519 | SKT**VLTGTKDTVCSGVTGAMNVAKGTIQTG**VDT |
| 326 | r8 | 100 | 87.9 | 0.519 | SKT**VLTGTKDTVCSGVTGAMNVAKGTIQTG**VDT |
| 359 | r9 | 75.8 | 81.8 | 0.569 | TKTVLTGTK**NTVCSGVTGAVNLA**KEAIQGGLDT |
| 392 | r10 | 57.6 | 63.6 | 0.635 | TKS**MVMGTKDTMSTGLTGAANVAKGAMQTG**LNT |
| 425 | r11 | 78.8 | 72.7 | 0.503 | TQN**IATGTKDTVCSGVTGAMNLARGTIQTG**VDT |
| 458 | r12 | 78.8 | 87.9 | 0.569 | TKI**VLTGTKDTVCSGVTGAANVAKGAVQGG**LDT |
| 491 | r13 | 75.8 | 81.8 | 0.519 | TKS**VLTGTKDAVSTGLTGAVNVAKGTVQTG**VDT |
| 524 | r14 | 78.8 | 90.9 | 0.569 | TKT**VLTGTKDTVCSGVTSAVNVAKGAVQGG**LDT |
| 557 | r15 | 66.7 | 75.8 | 0.635 | TKS**VVIGTKDTMSTGLTGAANVAKGAVQTG**VDT |
| 590 | r16 | 75.8 | 78.8 | 0.519 | AKT**VLTGTKDTVTTGLVGAVNVAKGTVQTG**MDT |
| 623 | r17 | 72.7 | 84.8 | 0.569 | TKT**VLTGTKDTIYSGVTSAVNVAKGAVQTG**LKT |
| 656 | r18 | 63.6 | 72.7 | 0.616 | T**QNIATGTKNTFGSGVTSAVNVAKGAAQTGV**DT |
| 689 | r19 | 75.8 | 78.8 | 0.519 | AKT**VLTGTKDTVTTGLMGAVNVAKGTVQTS**VDT |
| 722 | r20 | 84.8 | 87.9 | 0.569 | TKT**VLTGTKDTVCSGVTGAANVAKGAIQGG**LDT |
| 755 | r21 | 66.7 | 72.7 | 0.488 | TKS**VLTGTKDAVSTGLTGAVKLAKGTVQTG**MDT |
| 788 | r22 | 81.8 | 90.9 | 0.569 | TKT**VLTGTKDAVCSGVTGAANVAKGAVQMG**VDT |
| 821 | r23 | 81.8 | 87.9 | 0.569 | AKT**VLTGTKDTVCSGVTGAANVAKGAVQTG**LKT |
| 854 | r24 | 57.6 | 66.7 | 0.514 | TQNIATGTK**NTLGSGVTGAAKV**AKGAVQGGLDT |
| 887 | r25 | 72.7 | 78.8 | 0.519 | TKS**VLTGTKDAVSTGLTGAVNLAKGTVQTG**VDT |
| 920 | r26 | 93.9 | 93.9 | 0.519 | SKT**VLTGTKDTVCSGVTGAVNVAKGTVQTG**VDT |
| 953 | r27 | 72.7 | 75.8 | 0.552 | AKT**VLSGAKDAVTTGVTGAVNVAKGTVQTG**VDA |
| 986 | r28 | 72.7 | 72.7 | 0.635 | SKA**VLMGTKDTVFSGVTGAMSMAKGAVQGG**LDT |
| 1019 | r29 | 57.6 | 63.6 | 0.507 | TKTVLTGTKDA**VSAGLMGSGNVATGATH**TGLST |
| Consensus seq. | | 87.9 | 100 | n.d. | TKTVLTGTKDTVCSGVTGAVNVAKGAVQTGVDT |

(a) Position of the first residue according to Uniprot entry Q96Q06
(b) % sequence identity to repeat r6 (multiplied in patients) was determined
(c) % sequence identity to consensus sequence, determined using Weblogo
(d) Amyloidogenic propensity score for the individual 33-aa repeat was determined using ArchCandy
(e) AR within each repeat, as determined by ArchCandy, is shown in bold.

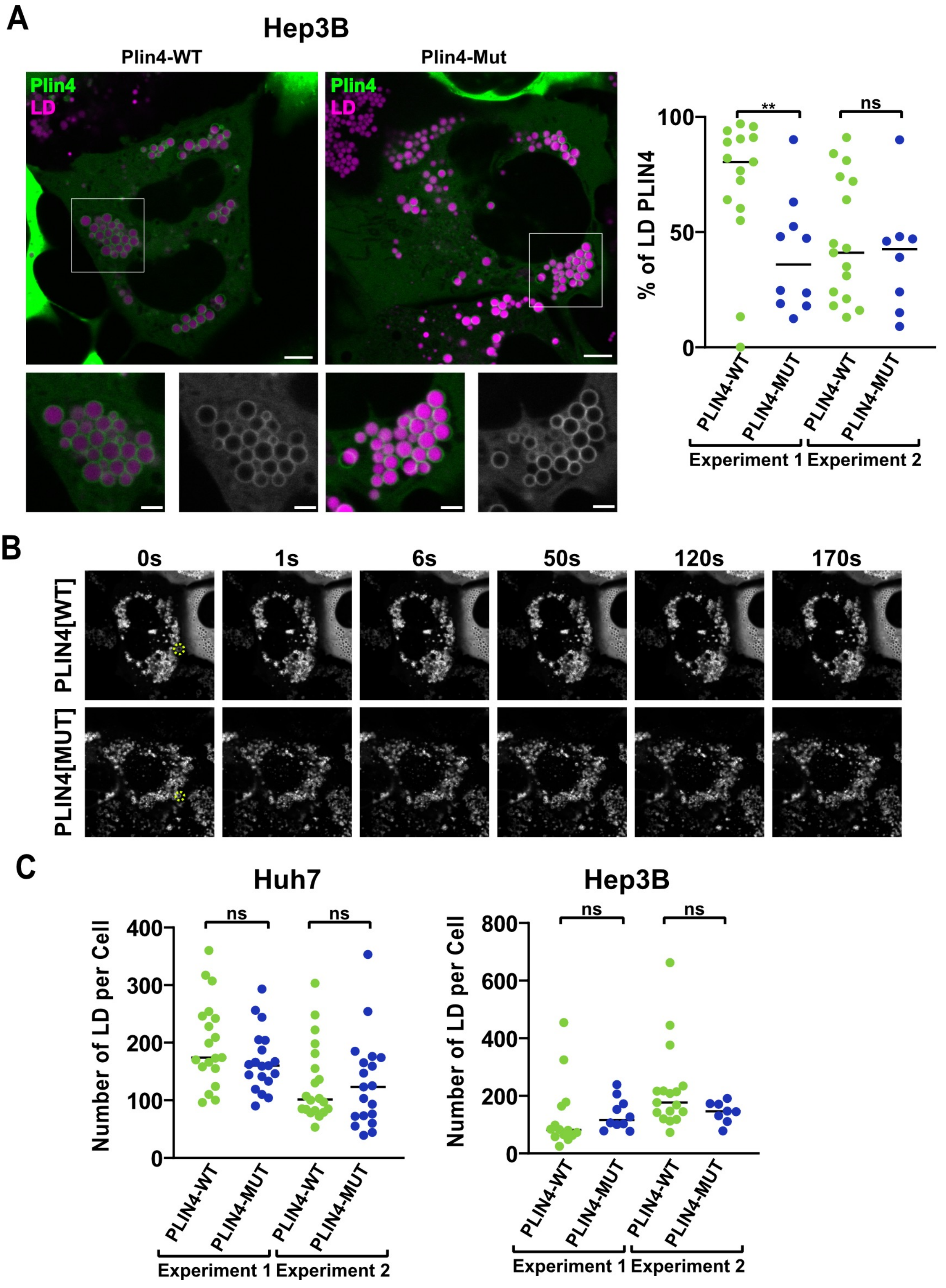


**Figure S1 (Related to Fig. 1). Comparison of the interaction of PLIN4-WT and PLIN4-Mut with LDs in cells. (A)** Localization of WT and Mut PLIN4-GFP in Hep3B cells after Dox induction in media containing oleic acid. LDs were stained with Lipi-Blue prior to live imaging by confocal microscopy. Bottom panels show a zoom-in of the selected areas. Scale bars: 5 µm and 2 µm for the zoom-in. Graphs show percent of LDs positive for PLIN4 in individual cells quantified in two independent experiments. Samples in the same experiment were compared using Mann-Whitney test, ns, nonsignificant (P>0.05)**,** **, P <0.01. **(B)** Examples of two FRAP experiments in Huh7 cells expressing PLIN4-WT-GFP (upper panels) or PLIN4-Mut-GFP (lower panels), used for quantification in Fig. S1E. Area that was bleached at t=0s is marked in yellow, and representitive images from the GFP taken during the recovery time-course are shown. **(C)** Quantification of the number of LDs per single z-section per cell in Huh7 cells (Fig. 1D) or Hep3B cells (Fig. S1A) expressing PLIN4-WT-GFP or PLIN4-Mut-GFP, from two independent experiments. Samples in the same experiment were compared using Mann-Whitney test, ns, nonsignificant (P>0.05).

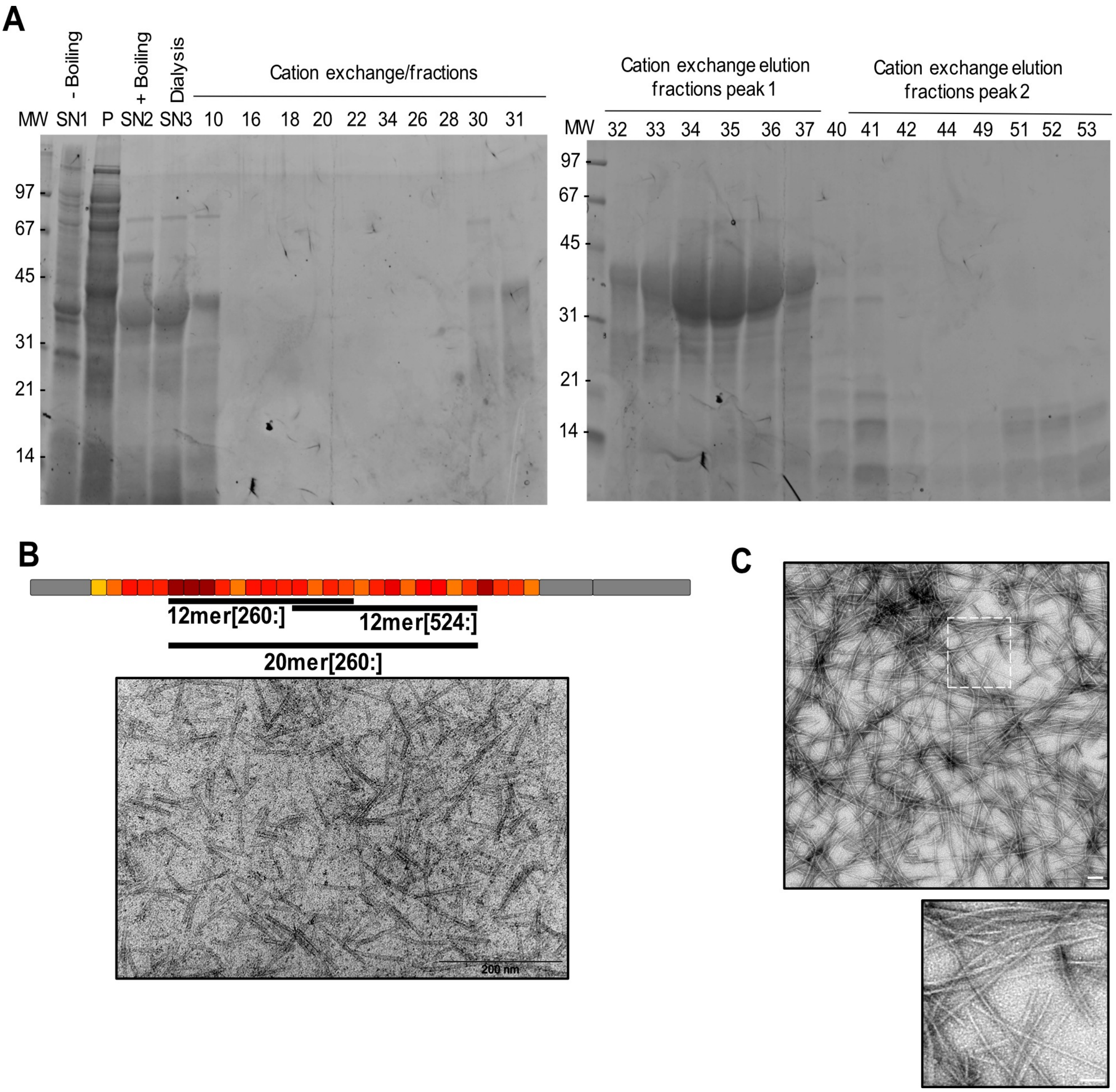


**Figure S2 (Related to Fig. 2). Purification of PLIN4-12mer[260:] and analysis of 20mer and 12mer[524:] by negative-stain EM. (A)** Purification of 12mer[260:]. Lanes are labeled as follows: MW, molecular weight ladder; P, pellet after centrifugation; SN1, soluble fraction after centrifugation; SN2 ("+boiling"), heat-resistant fraction obtained by boiling SN1; SN3, dialyzed SN2, prepared for cation exchange chromatography. The dialyzed fraction (SN3) was subjected to cation exchange chromatography using a NaCl gradient (1 mM to 1000 mM). Elution fractions were collected and analyzed: Fractions 10–28 correspond to flow-through, fractions 30–37 contain the major elution peak with intact protein form, and fractions 40–53 represent degraded protein form. Samples were analyzed on 12% SDS-PAGE and stained with Sypro Orange. Expected molecular weight of PLIN4-12mer[260:] is 38.1 kDa. **(B)** Negative stain EM of PLIN4-20mer fibrils, obtained after prolonged incubation (>1 week) of purified protein (1.5 mg/ml, see ref. 18) in HK buffer containing 1 mM MgCl2 and 1 mM DTT at room temp. A diagram of PLIN4 depicting the position of the PLIN4-20mer fragment compared to 12mer[260:] and 12mer[524:] is shown above the image. **(C)** Negative stain EM of fibrils obtained with purified 12mer[524:] (2 mg/ml) after prolonged incubation (>1 week) with shaking at 37°C. A 4x zoom-in of the marked area is shown on the right. Scale bars: 200 nm / 50 nm for the zoom-in.

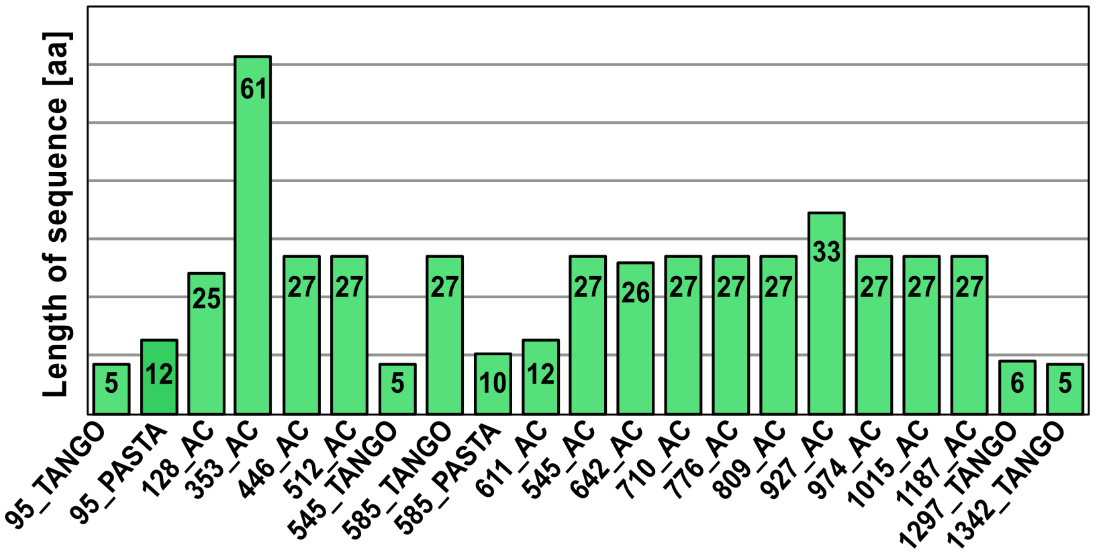


**Figure S3 (Related to Fig. 4). Analysis of amyloidogenic properties of PLIN4 using the TAPASS pipeline.** The plot represents the length (in aa) and position of all ARs identified within PLIN4. The position of the first aa of each AR and the algorithm that identified it (TANGO, PASTA or ArchCandy – AC) is indicated on the x-axis. The repetitive region (aa95-1051) is indicated under the x-axis. See also Table S1.

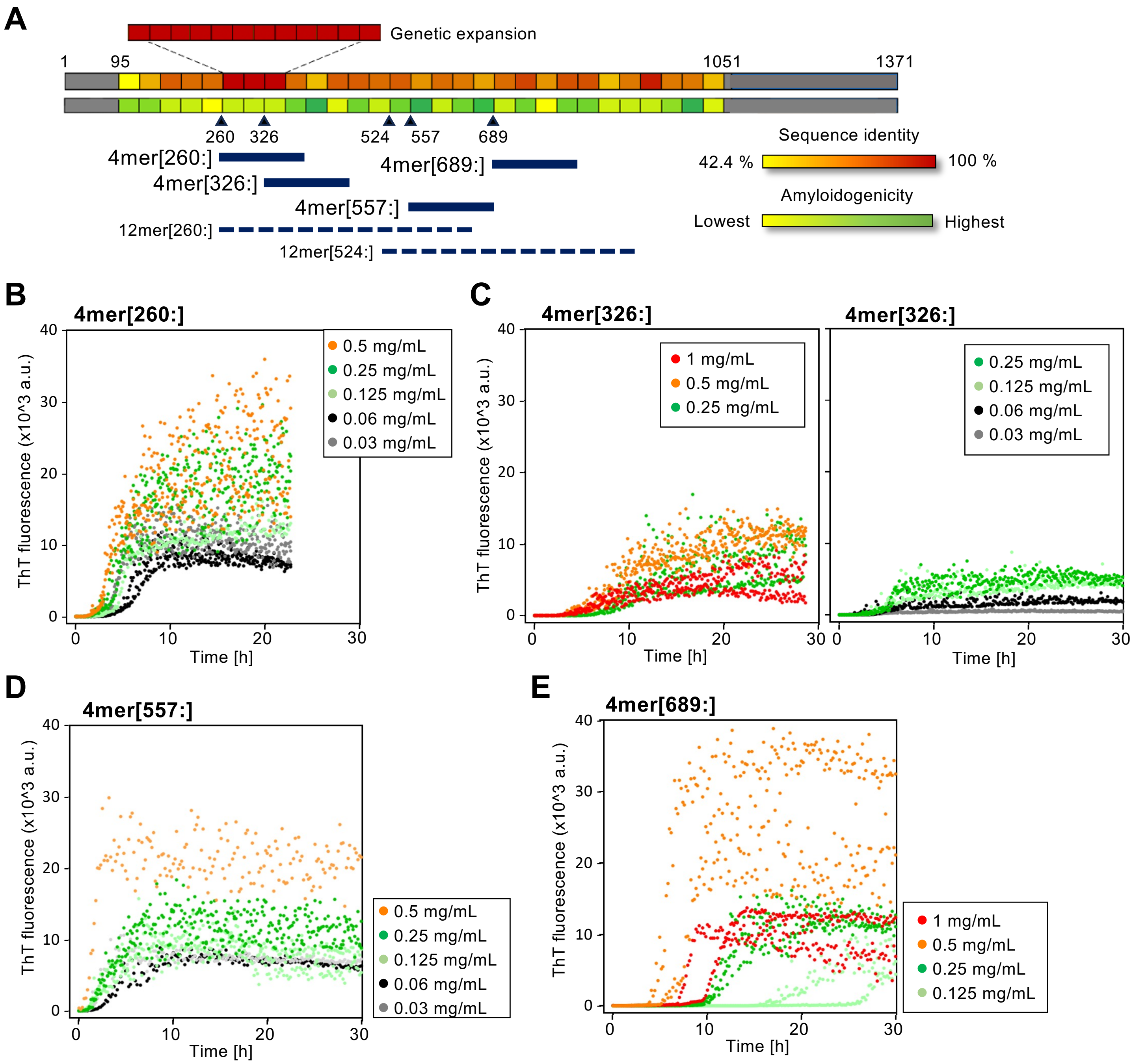


**Figure S4 (Related to Fig. 4 and Table S1). Analysis of the aggregation kinetics of PLIN4 4mer constructs. (A)** Diagram of PLIN4 showing % identity between individual 33-aa repeats and their amyloidogenic propensity (as in Figure 3). The position of PLIN4 4mer's is indicated under the diagrams. All 4mer constructs map within the two PLIN4 12mer's, [260:] and [524:], indicated with dashed lines. **(B)** Aggregation of 4mer[260:] in the ThT assay is fast even at lower protein concentrations, down to 0.03 mg/ml, and shows little concentration dependence (compare to Fig. 4C). **(C)** Aggregation of 4mer[326:], which contains only one of the three identical repeats present in 4mer[260:], is slower than that of 4mer[260:], and requires a higher protein concentration. Note that this construct also contains one repeat with the highest amyloidogenic score (r10, 0.635), followed by a repeat with a low score (r11, 0.503). The difference in aggregation kinetics of 4mer[260:] and 4mer[326:] is therefore likely due to a combination of several factors. For clarity, aggregation curves from the same experiment are presented in two different graphs. **(D)** 4mer[557:], which contains two repeats with high amyloidogenic scores (r15, 0.635 & r18, 0.616), but no identical repeats, aggregates at a similar rate and protein concentration as 4mer[260:]. **(E)** 4mer[689:], which contains the repeat 1mer[755:] with the lowest amyloidogenic score (r21, 0.488) and no highly amyloidogenic repeats shows a longer lag-phase and requires a higher protein concentration for robust aggregation. Aggregation of this construct was more variable between independent experiments.

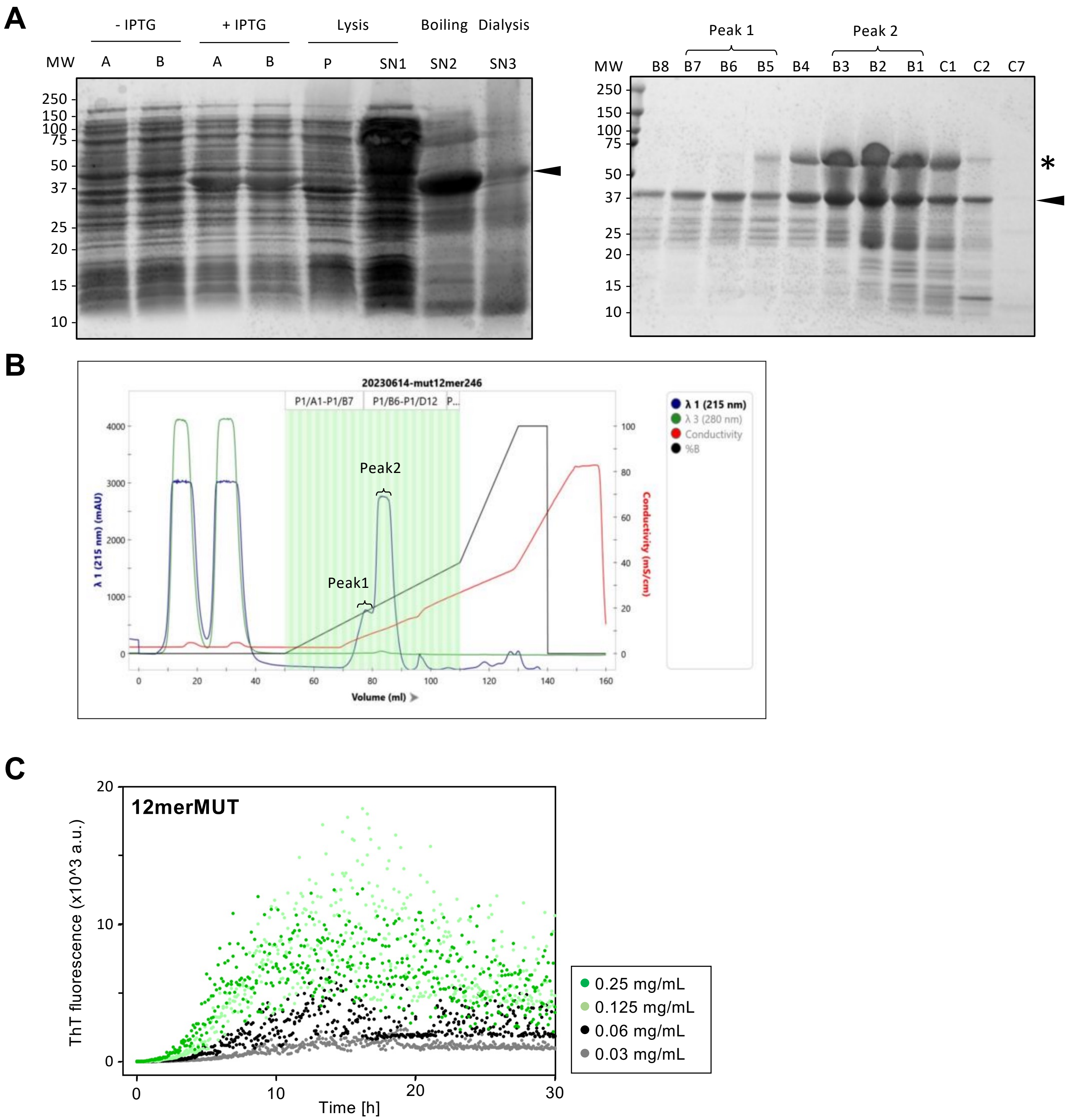


**Figure S4 (Related to Fig. 4). Purification and characterization of PLIN4-12merMUT. (A)** Purification of PLIN4-12merMUT was achieved following the same procedure as for 12mer[260:] (Fig. S2A). Fractions were analyzed by SDS-PAGE and stained with Coomassie Blue. The right gel shows fractions from the cation exchange chromatography of the supernatant (SN3) after dialysis. Two peaks were analyzed, the chromatogram is shown in panel **(B)**. Arrows indicate the predicted size of 12merMUT (~37 kDa). (*) indicates oligomers of 12merMUT, which were removed by centrifugation before aggregation assays. **(C)** ThT aggregation assay with purified 12merMUT at lower concentrations compared to the assay shown in Fig. 4D, right panel. Some aggregation is observed at concentrations as low as 0.06 mg/ml.

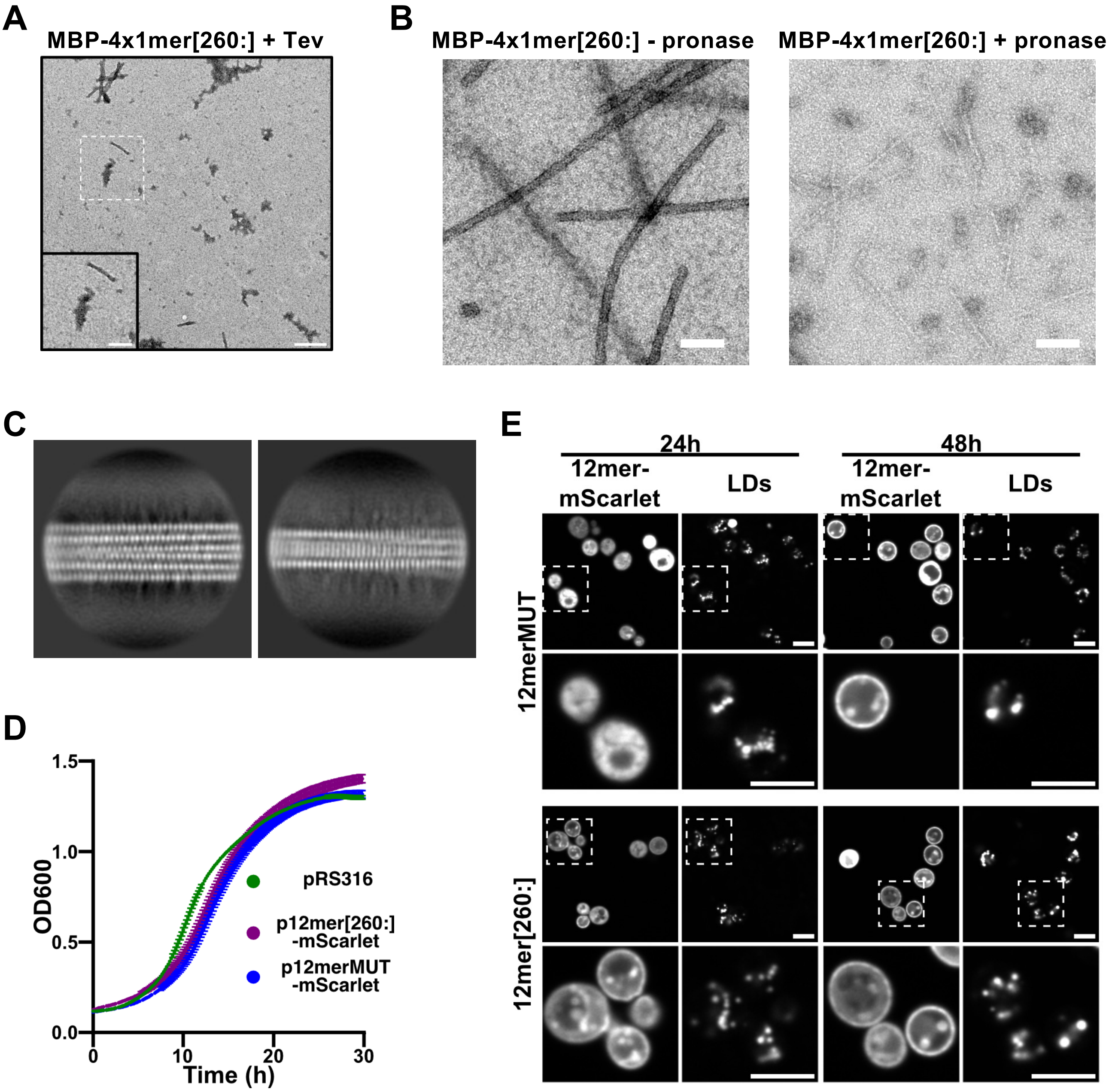


**Figure S5 (Related to Fig. 4). Additional analysis of fibrils formed by MBP-4x1mer[260:] and of yeast expressing PLIN4-12mer constructs.** **(A)** Reaction mixtures of MBP constructs shown in Fig. 4F after ThT aggregation were analyzed by negative stain EM. Fibrils were detected only in the MBP-4x1mer[260:] samples -TEV (shown here, conc = 1.1 mg/ml) and +TEV (Fig. 4G). Scale bar: 1 µm or 0.5 µm for the magnified area (2 x zoom). **(B)** Filaments formed by MBP-4x1mer[260:] (2 mg/ml, no Tev treatment) at 37°C were observed by negative stain EM, revealing thickness of 20 nm and length of several microns (left panel, no pronase treatment). The same filaments after 25 min treatment with pronase (0.5 mg/ml at 20°C) display a reduced thickness ranging from 5 to 8 nm. Scale bar: 100 nm. **(C)** Representative 2D classes following the processing of cryo-EM images of untreated MBP-4x1mer[260:] filament. 2D classification shows classes very similar to those obtained with PLIN4-12mer[260:] for thick (left panel) and thin (right panel) polymorphs. **(D)** Expression of PLIN4-12mer constructs in budding yeast does not affect cell growth**.** Growth of yeast strains containing either an empty plasmid (pRS316), or a plasmid expressing the indicated construct, 12merMUT-mScarlet and 12mer[260:]-mScarlet, from a constitutive (GPD) promotor. **(E)** Localization of 12merMUT-mScarlet (upper panels) and 12mer[260:]-mScarlet (lower panels) in aged yeast cells that accumulate LDs (stained with AutoDOT), after 24h and 48h incubation at 30°C, as indicated. Bottom rows for each construct show a 3x zoom-in of the indicated area. Scale bar: 5 µm.

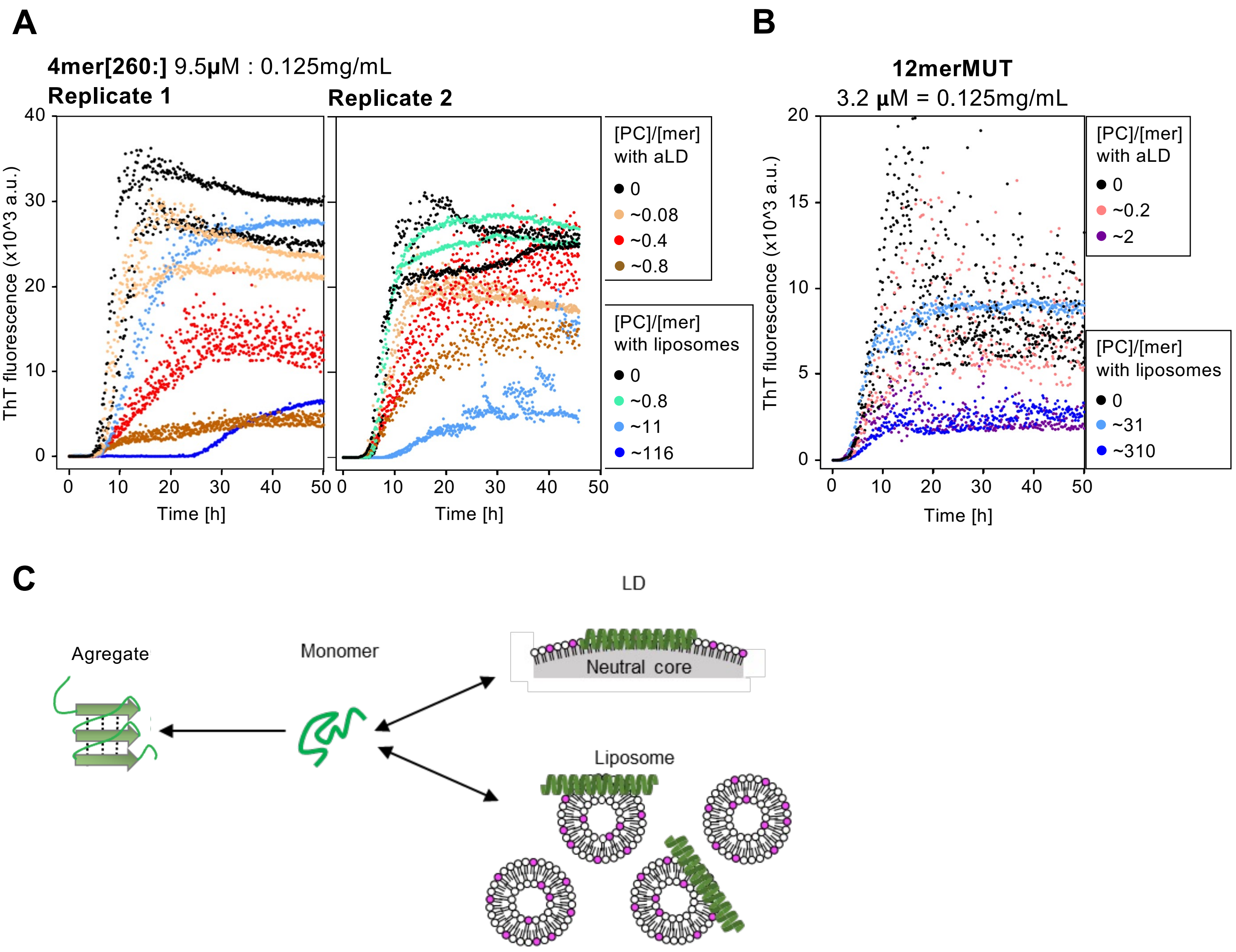


**Figure S7 (Related to Fig. 5). Aggregation of PLIN4 repetitive fragments in the presence of liposomes. (A, B)** The ThT aggregation assay of 4mer[260:] at 0.125 mg/mL (9.5 µM) (A), or 12merMUT at 0.125 mg/mL (3.2 µM) (B), in buffer (black symbols), in the presence of artificial LDs (aLDs) as in Fig. 5C (brown, red or pink symbols, indicating the PC to repeat (mer) ratio), or in the presence of diphytanoyl-PC liposomes (green or blue symbols, indicating the PC to repeat (mer) ratio). LDs were ~40 times larger in diameter than liposomes, as shown in (C). Note that due to the LD and liposome preparation protocols, the LD concentration in the aggregation reactions cannot be higher than the maximum concentration indicated in the panels. For 4mer[260:] (A), two independent experiments are shown (Replicate 1 & Replicate 2), with some variability between experiments. Replicate 1 represents the same experiment as shown in Fig. 5C (left panel). Note that LDs influence PLIN4 aggregation at a lower PC concentration than liposomes, consistent with a higher affinity of the PLIN4 amphipathic helix for LDs than for liposomes. For 12mer-MUT (B), the experiment shown is an independent experiment from that shown in Fig. 5C. **(C)** Schematic depicting the repetitive region of PLIN4, which is disordered in solution and can fold into an amphipathic helix in contact with dipthytnaoyl liposomes or LDs, or form amyloid fibrils.
